# scDALI: Modelling allelic heterogeneity of DNA accessibility in single-cells reveals context-specific genetic regulation

**DOI:** 10.1101/2021.03.19.436142

**Authors:** T. Heinen, S. Secchia, J. Reddington, B. Zhao, E.E.M. Furlong, O. Stegle

## Abstract

While the functional impact of genetic variation can vary across cell types and states, capturing this diversity remains challenging. Current studies, using bulk sequencing, ignore much of this heterogeneity, reducing discovery and explanatory power. Single-cell approaches combined with F1 genetic designs provide a new opportunity to address this problem, however suitable computational methods to model these complex relationships are lacking.

Here, we developed scDALI, an analysis framework that integrates single-cell chromatin accessibility for unbiased cell state identification with allelic quantifications to assay genetic effects. scDALI builds on Gaussian process regression and can differentiate between homogeneous (pervasive) allelic imbalances and cell state-specific regulation. As a proof-of-principle, we applied scDALI to whole *Drosophila* embryos from F1 crosses, profiling sciATAC-seq at three embryonic stages. Even in these very complex samples, scDALI discovered hundreds of peaks with heterogeneous allelic imbalance, having effects in specific lineages and/or developmental stages. Our study provides a general strategy to identify the cellular context of allelic imbalance, a crucial step in linking genetic traits to cellular phenotypes.

## Introduction

The functional impact of genetic variants on molecular traits such as gene expression can be influenced by the cell type or cell state. Particularly non-coding variants in enhancer elements can impact a gene’s expression in one tissue and not in others. Population-scale genetics studies, using bulk sequencing across individuals, have identified many such tissue-specific^1–3^ and developmental stage-specific^4^ effects, which often involve rare genetic variants. However, even dissected tissues are composed of heterogeneous cell types, thus motivating the application of single-cell sequencing to reveal cell state dependencies of genetic effects. Recent single-cell studies in *in vitro* models revealed changing genetic dependencies across different cellular transitions^5,6^. However, such population-scale *in vitro* models do not fully recapitulate the complexity of a tissue’s or embryo’s development, and additionally require large numbers of assayed individuals, which limits their applicability, particularly *in vivo*. The latter can be addressed by measuring allele-specific signals, i.e., quantifying molecular traits separately for each haplotype^7–10^, which can identify genetic effects even in a single individual. Integrating both approaches, by quantifying allelic imbalance in single cell data, could be a powerful strategy to dissect the functional impact of genetic variants both within and across cell types simultaneously. Prior studies have measured allele-specific properties at a single-cell level to characterize transcriptional bursting and stochasticity in gene expression^11,12^. However, single-cell studies that consider allelic regulation are only beginning to emerge^13,14^, and computational methods to model these complex relationships are not established.

To address this, we developed a generalizable computational model and analysis framework, scDALI (**s**ingle-**c**ell **d**ifferential **a**llelic **i**mbalance), that leverages allele-specific analysis of single-cell data to decode context-specific genetic regulation. We applied this approach to study allele-specific variation in single cell chromatin accessibility data (sciATAC-seq) in F1 embryos at three stages of embryogenesis, generated from a cross of four isogenic *Drosophila melanogaster* strains. The model identified hundreds of regulatory regions with allelic imbalances in specific cell types or developmental stages. Among these effects we identify regions, putative enhancers, with opposing allelic imbalance in different cell lineages, which are missed by bulk assay profiling. Obtaining equivalent information with current strategies would require substantial cell sorting, which in addition to being laborious, requires cell markers that are not always available for cells in specific transitions. scDALI is a widely applicable computational method that leverages single-cell technologies to avoid cell sorting, providing the means to discover and quantify the functional impact of cell state-specific genetic effects in an unbiased manner.

## Results

scDALI combines genetic information on allelic imbalance, either observed in outbred individuals or generated from F1 crosses of inbred wild-isolates, with single-cell chromatin accessibility profiling, to chart the cell state dependency of genetic regulation. Key to our approach is the integration of two independent signals that can be obtained from the same single-cell ATAC-seq (scATAC-seq) experiment: total accessibility counts and allele-specific quantifications. scDALI first uses total accessibility, quantified at ATAC-seq peaks, to define cell types and cell states, similar to established workflows for the inference of state clusters^15^ or pseudo-temporal orderings^16^. Second, from the same dataset, allele-specific accessibility counts from matched cells are extracted, which allow for quantifying allelic imbalances and therefore genetic effects (**Fig. 1a**).

**Figure 1.**
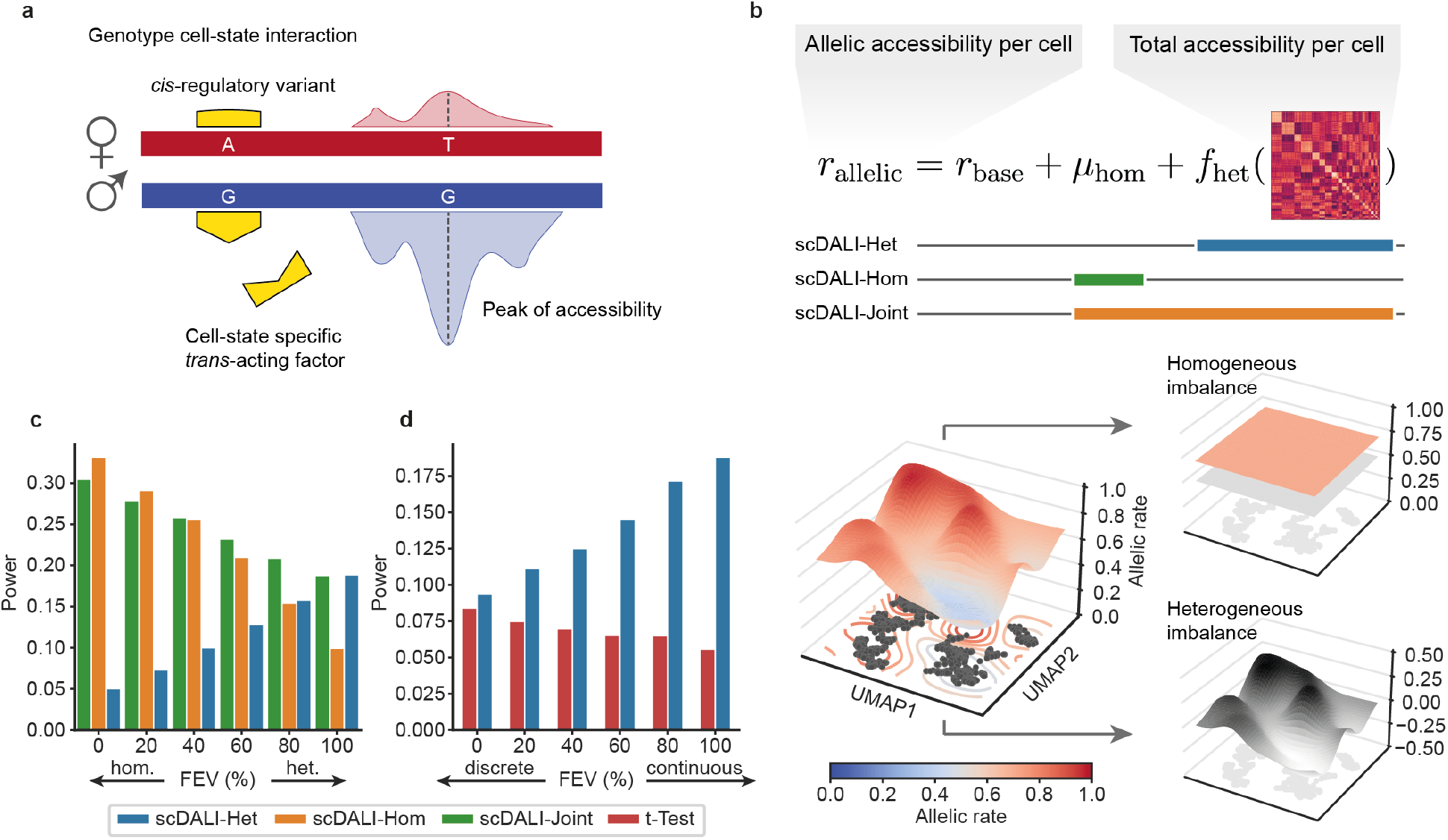
scDALI overview and model validation using simulated data. **(a)** Causes and quantification of allelic imbalance from sciATAC-seq data. Heterozygous variants within peaks of accessibility (T and C variant) are used to assign reads to either haplotype. A *cis*-regulatory genetic variant (A and G variant, yellow) impacts the efficacy of a *trans*-acting chromatin regulator on the maternal allele, which results in allelic imbalance. If the *trans*-acting factor is cell state-specific, this effect will be heterogeneous across cell states, otherwise homogeneous. **(b)** Integration of total accessibility and allelic counts per cell. scDALI models allelic imbalance in single cells as the sum of a fixed expected rate (*r*_*base*_, e.g., 0.5 for autosomes), a global homogeneous component (*μ*_*hom*_) and a heterogeneous cell state-specific component (*f*_*het*_), which is characterized by a cell state kernel matrix. Three alternative score tests allow for assessing the evidence for either global allelic imbalance (scDALI-Hom), heterogeneity across cell states (scDALI-Het) or to jointly test for either form of allelic imbalance (scDALI-Joint). **(c-d)** Assessment of power of alternative tests for allelic imbalance using simulated data. Allelic counts were simulated using different cell state kernels derived from real data (**Supp. Fig. 3, Methods**). **(c)** Varying the simulated fraction of explained variance (FEV) of heterogeneous vs. homogeneous allelic imbalance. Whereas scDALI-Het and scDALI-Hom detect signals with simulated persistent or heterogeneous effects respectively, scDALI-Joint identifies either type of allelic imbalance. **(d)** Varying between simulating allelic effects from a discrete and continuous cell state model. scDALI-Het is better powered than a baseline approach based on a t-test to perform cluster-wise one-vs-all tests (25 clusters; Bonferroni adjusted for the number of clusters; **Methods**).

scDALI is a probabilistic model that connects both of these signals, while aggregating evidence across cells to mitigate the sparsity of single cell data. The model can be used to estimate the landscape of allelic rates across cell states, and to comprehensively test for sites that exhibit different types of allelic imbalance. In particular, the model allows for differentiating between homogeneous (pervasive) allelic imbalance across all cells and heterogeneous effects that vary across cell types and cell states (**Fig. 1b**). Briefly, our method is based on Gaussian process (GP) regression with a Beta-Binomial likelihood to capture count noise and residual overdispersion due to unmodelled variability in the data (**Methods**). This formulation extends the classical Beta-Binomial framework that is commonly used for the allele-specific analysis of bulk sequencing data^7,9,10,17^. In addition to capturing homogeneous deviation from a specified base allelic rate (e.g., 0.5 for autosomes in diploid organisms), scDALI incorporates a cell state kernel matrix to account for heterogeneity and to assess its effect on allele-specific variation. Similar to previous applications of GPs to single-cell RNA-seq^18,19^, the GP framework provides the flexibility to encode a variety of cell state effects, including discrete cell clusters as well as continuous developmental trajectories. Within the scDALI framework, we formulate computationally efficient score tests^20,21^ to identify different types of allelic imbalance: Global homogeneous imbalance (scDALI-Hom), heterogeneity across cell states (scDALI-Het) or either of these effects (scDALI-Joint) (**Fig. 1b**). Additionally, the model estimates what fraction of the total allele-specific variance can be explained by cell state effects and can be used to estimate and visualize allelic imbalances across cell states (**Methods**).

While in principle any method for cell state inference can be used in conjunction with scDALI, the software implementation comes with a workflow for cell state inference based on a variational autoencoder (VAE)^22^. VAEs have previously been proposed for scRNA-seq data^23^, and can similarly be adapted to scATAC-seq^24^. A VAE is a neural network with a probabilistic bottleneck layer that learns the distribution of the data by compressing high-dimensional observations into a lower dimensional latent space. Our implementation (**Supp. Fig. 1a)** not only integrates scATAC-seq profiles across datasets and batches, but also allows to explicitly model information about different sampling times for developmental datasets (**Supp. Fig. 1d, e**). This extension of previous VAE formulations enables our model to infer continuous temporal ordering of cells by coupling the VAE objective function with a regression problem to predict sampling time from the latent cell state representation (**Methods**).

### Model validation using simulated data

Initially we validated our approach using simulated data, which was designed to mimic real data by adapting key parameters from empirical sciATAC-seq profiles from developing *Drosophila melanogaster* embryos (cell states, overdispersion parameters; **Supp. Fig. 2a, Methods**). First, we assessed the calibration of all three scDALI tests by simulating from the corresponding null models of each test, confirming uniformly distributed p-values (**Supp. Fig. 2b-e**). Next, we simulated allelic counts from the alternative model, varying the proportion of homogeneous versus heterogeneous allelic imbalance (**Fig. 1c, Supp. Fig. 3, 4a**). The results demonstrate that scDALI-Joint identifies effects of both classes, generalizing the individual tests scDALI-Het and scDALI-Hom. We then went on to simulate allelic counts by drawing from scDALI-Het, considering either a kernel that reflects 25 discrete clusters, continuous states or weighted combinations thereof (**Fig. 1d, Supp. Fig. 3, 4b**). When comparing scDALI-Het with a cluster-wise one-vs-all t-test (Bonferroni adjusted for the number of clusters; **Methods**), we observed that scDALI-Het was noticeably better powered than this baseline. We considered a range of additional settings, varying the levels of overdispersion and kernel variance, consistently observing that scDALI was better powered than alternative strategies to identify allelic imbalance (**Supp. Fig. 4**). scDALI is implemented as open-source software and computationally efficient, scaling to the analysis of large datasets with up to tens of thousands of cells (**Supp. Fig. 5**).

### scDALI identifies heterogeneous allelic imbalance in developing *Drosophila* embryos

Next, we applied scDALI to study context-specific regulation across embryonic development in *Drosophila melanogaster*. We profiled single cell chromatin accessibility by sciATAC-seq in F1 hybrid embryos obtained by mating the same mother with four genetically distinct fathers^17^. To ensure that we capture regulatory variation associated with major developmental events, we profiled embryos from the four F1 crosses at three key stages of embryonic development (**Fig. 2a**, 2-4 hours, 6-8 hours and 10-12 hours after egg laying), which correspond to stages when the majority of cells are multipotent, or are undergoing lineage commitment, and or tissue differentiation respectively. Sequencing the resulting 12 sciATAC-seq libraries generated a dataset of 35,485 single cells (between 8,000 and 10,000 cells per cross) that passed stringent quality metrics (**Supp. Fig. 6**; **Methods**). Overall, our dataset has all the hallmarks of high quality sciATAC-seq, including the appropriate nuclesomeal banding pattern (**Supp. Fig. 6a**), and a high concordance to previously identified peaks from a time-matched sciATAC-seq dataset in a reference strain^25^ (**Supp. Fig. 7**). The top 25,000 most accessible of these peaks across all crosses and time points were used for the definition of cell states.

**Figure 2.**
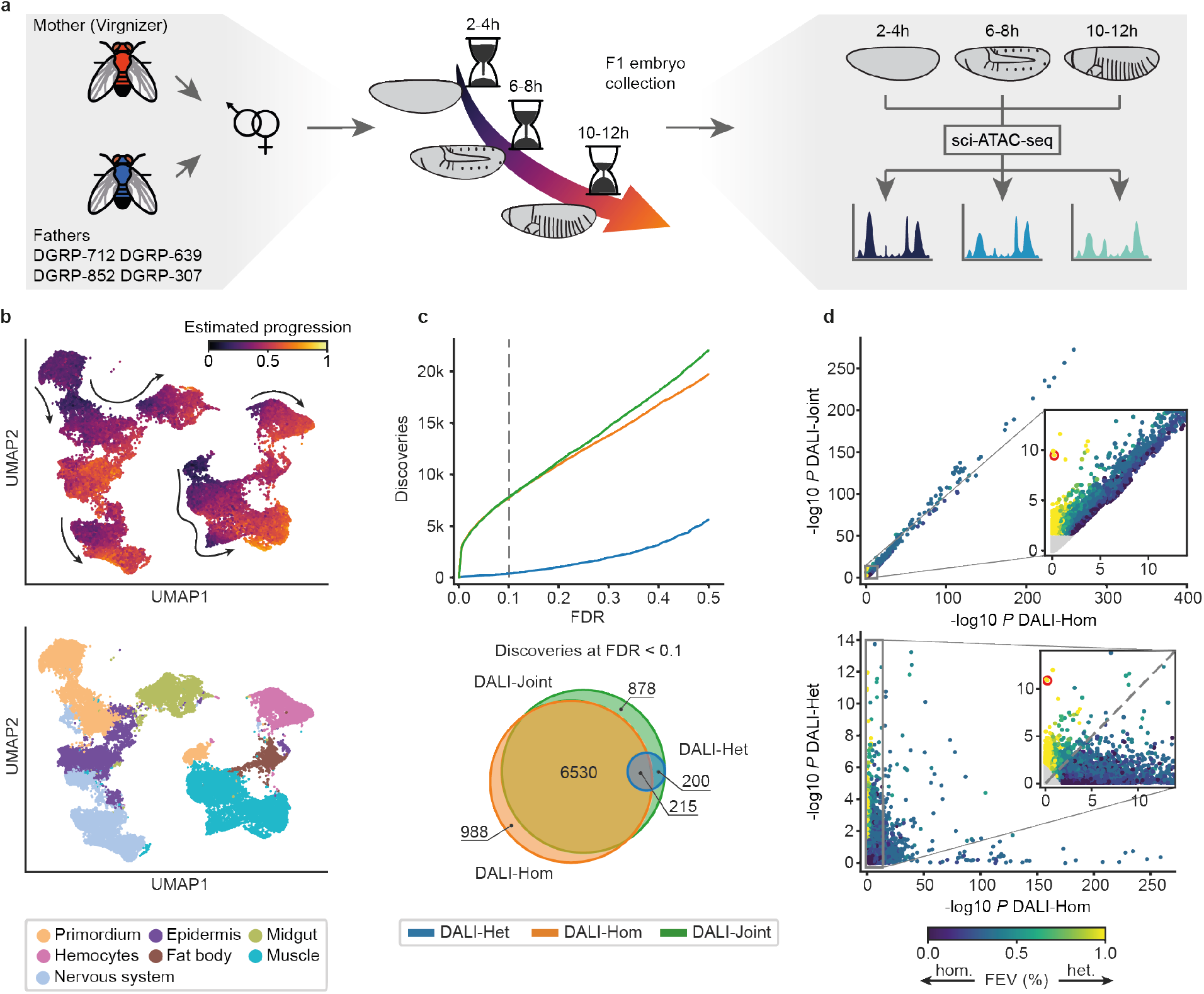
Application of scDALI to sciATAC-seq of Drosophila F1 embryos. (**a**) Experimental design. Chromatin accessibility was profiled in four F1 crosses at three distinct developmental stages (2-4, 6-8 and 10-12 hours after egg laying), resulting in 12 sciATAC-seq libraries. (**b**) UMAP visualization of the full dataset (34,053 cells from 12 sciATAC-libraries, excluding cell clusters with ambiguous annotations) based on the latent space of the Variational Autoencoder (VAE) (**Methods**). Top: Cells colours by the continuous temporal ordering as estimated from the VAE model. Bottom: Coloured by the major lineage annotation. **(c)** Number of sites across crosses and time points with allelic imbalance identified by scDALI-Joint, scDALI-Hom and scDALI-Het. Top: Number of discoveries as a function of the FDR (Benjamini Hochberg adjusted). Bottom: Overlap between the sites identified by all three scDALI tests (FDR < 0.1). (**d**) Scatter plot of negative log P-values between scDALI-Joint and scDALI-Hom (top) and scDALI-Het versus scDALI-Hom (bottom) respectively. Colour denotes the estimated fraction of allele-specific variance explained by cell state effects; points in grey are not significant (adjusted scDALI-Joint *P* > 0.1). Inset plots zoom in on peaks with pronounced cell state-specific imbalances. The red circle highlights the peak chr3R:20310056-20311056, a region showing prominent cell state-specific effects with no discernable global imbalance (c.f. **Fig 3a-c**).

To define a common cell state space across the full dataset, we applied the VAE cell state inference workflow that is part of scDALI (**Methods**). The VAE model inferred a common latent space for all F1 crosses in which the cells do not cluster by sample (**Supp. Fig. 1c**), but rather they arrange progressively by developmental time (**Fig. 2b, Supp. Fig. 1d, e**). Cells were then clustered based on this lower-dimensional representation using the Leiden algorithm^26,27^ (28 clusters, **Supp. Fig. 1b)**. Cell type identities were assigned to each cluster based on their enrichments for open-chromatin spanning cell type-specific enhancers, using a large compendium of enhancers with curated spatio-temporal activity during embryogenesis and genes with known tissue-specific expression^25^ (**Methods**). Four smaller clusters with ambiguous annotations that likely correspond to barcode collisions were excluded from further analysis. This annotation process resolved seven cell populations that are representative of major embryonic lineages, including muscle, nervous system and ectoderm (**Fig. 2b**).

Next, we quantified chromatin accessibility on an allele-specific level. To avoid allelic mapping artifacts, we applied WASP ^10^, filtering between 7-8% of mapped reads (**Supp. Fig. 8a, Methods**). Allele-specific chromatin accessibility was quantified within 1kb regions centered on ATAC peaks, requiring that each read overlapped at least one heterozygous variant. This resulted in a haplotype assignment for 20% of the reads (based on 5-6 variants per region on average, **Supp. Fig. 8b, c**). After discarding peaks with low allelic coverage (mean count of reads that could be assigned to either allele < 0.1), we obtained between 8,040 and 12,861 peaks per cross for further analysis resulting in a combined set of 39,530 peaks to be tested (**Supp. Fig. 8d, e**).

We applied scDALI to test for and characterize allelic imbalance for all crosses and timepoints (**Fig. 2c, d, Supp. Fig. 9**), using the VAE latent space coordinates as cell state representations. scDALI-Joint identified 7,823 out of 39,530 peaks with evidence for allelic imbalance (FDR < 0.1, Benjamini-Hochberg adjusted). Notably, the majority of these peaks were also identified by scDALI-Hom (83%), indicating that homogeneous imbalance is prevalent. However, scDALI-Het identified 415 sites with evidence for cell state-specific allelic imbalance, all of which were also detected by scDALI-Joint. Notably, 200 sites detected by scDALI-Het were missed by scDALI-Hom, indicating that strong heterogeneity can preclude the identification of allelic imbalance by bulk sequencing or analysis strategies that assume exclusive homogenous effects. For instance, a peak (region chr3R:20310056-20311056) in cross F1-DGRP-307 was found to be highly significant by scDALI-Joint (*P* = 1.93×10^−8^) and scDALI-Het (*P* = 5.45×10^−8^) but was globally consistent with a model that assumes no allelic imbalance (scDALI-Hom *P* = 0.81, **Fig. 2d**).

### Properties of regions with heterogeneous allelic imbalance

To better understand the structure of allelic variation among the 415 heterogeneously imbalanced regions, we used scDALI to estimate allelic rates across the VAE cell state space for each peak (**Methods**). These estimates allowed us to visualize the landscape of allelic imbalance on top of a two-dimensional UMAP representation of the VAE space and to assess the magnitude and dispersion of allelic rates within and between different cell populations. While our analysis focused on the 7 major lineages, scDALI can in principle also be applied in an entirely unbiased manner if no such reference annotation is available.

We find several cases in which allelic imbalance affects known lineage-specific regulatory elements. For example, region chr3R:22877489-22878489 (scDALI-Het *P* = 2.7×10^−5^) has been previously identified as a neuronal-specific DNase Hypersensitive Site (DHS)^28^ and has been demonstrated to function as a nervous system enhancer *in vivo* (CAD4 database^25^). Accordingly, this region is identified as predominantly accessible in the nervous system (**Fig. 3a**). In addition, while cells from other lineages show no appreciable allelic imbalance, accessibility in the nervous system is strongly biased for the paternal allele (**Fig. 3b, c**). Ordering cells by their allelic rate and computing the difference between the top and bottom 10% quantiles (Qdiff10), we define a measure of the effect size of heterogeneous allele-specific imbalances, which captures the variation in allelic rates between the most extreme populations (**Fig. 3d**). In this example we obtain a Qdiff10 of 0.24 despite the overall mean allelic rate being close to 0.5 (scDALI-Hom *P* = 2.9×10^−3^).

**Figure 3.**
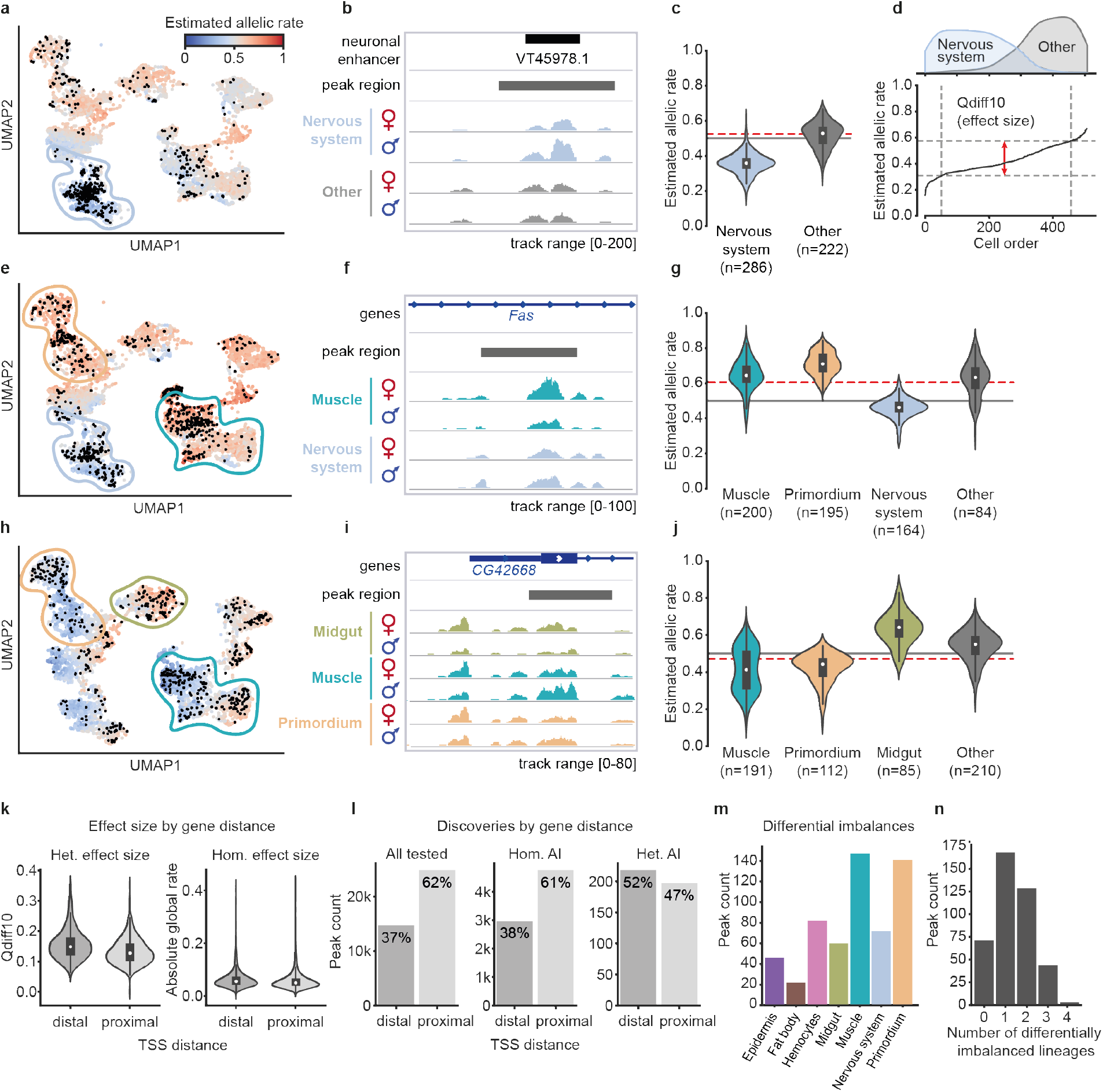
Examples and analysis of ATAC peaks with heterogeneous allelic imbalance. **(a)** UMAP visualization displaying cells colored by their estimated allelic rate (maternal accessibility relative to total accessibility) for region chr3R:22877489-22878489 in cross F1-DGRP-639. Black dots indicate cells with observed allelic rates (non-zero allelic total counts). **(b)** Genome browser tracks for region chr3R:22877489-22878489 illustrating allele-specific aggregate accessibility for the nervous system and other populations. **(c)** Distribution of estimated allelic rates stratified for cells from the nervous system vs. other populations. Red line shows the estimated overall allelic rate *μ*_*hom*_ across all cells. **(d)** Ordered estimated allelic rates. The difference between 10% and 90% quantiles is used as a measure of effect size for heterogeneous allelic imbalance (Qdiff10). **(e-g)** Example region chr2R:13675707-13676707 in cross F1-DGRP-639, revealing opposing effects in the nervous system and muscle lineage. **(h-j)** Example region chr3R:20310056-20311056 in F1-DGRP-307, revealing lineage-specific differences in allelic rates for the muscle, primordium and midgut, as well as intra-lineage variation within the muscle population. **(k)** Violin plots of effect size estimates for heterogeneously (10% quantile difference as in **d**) and homogeneously imbalanced (absolute deviation from .5) peaks, considering distal and promoter-proximal regions separately. Distal peaks are associated with both larger absolute allelic imbalance and stronger heterogeneity (*P* = 5.6×10-5, *P* = 2.17×10-26, one-sided Mann-Whitney U test). **(l)** Total number of peaks tested and peaks with allelic imbalance identified using alternative tests (FDR < 0.1), stratified by the peak distance to the transcription start site (TSS) of the closest gene. Heterogeneously imbalanced peaks are markedly more common at distal regions. **(m, n)** By-lineage analysis of allelic imbalance using scDALI-Het for peaks with significant heterogeneous imbalances. **(m)** Distribution of the number of differentially imbalanced lineages per peak. **(n)** Distribution of the number of peaks with increasing numbers of differentially imbalanced lineages. The majority of peaks show imbalance in a single lineage.

Interestingly, we find a number of regulatory regions that show opposing allelic imbalances in different lineages. For example, region chr2R:13675707-13676707, has only a small maternal bias (estimated overall mean 0.61) when considering the global allelic rate, but is identified as a site with pronounced allelic heterogeneity by scDALI (scDALI-Het *P* = 1.5×10^−8^, **Fig. 3e**). This region was previously identified as a neuronal and muscle-specific DHS^28^ and accordingly shows increased accessibility in the nervous system and muscle in our data. However, accessibility is biased for the maternal allele in the muscle and the paternal allele in the nervous system (Qdiff10 = 0.29, **Fig. 3f, g**). Another example is chr3R:20310056-20311056 (scDALI-Het *P* = 5.45×10^−8^), a region spanning an intron of the gene *CG42668*. The total accessibility of this region largely coincides with the known tissue-specific gene expression of *CG42668* in the cells of the primordium, midgut and visceral muscle. Our allele-specific analysis revealed differential allele-specific effects in all three cell types, suggesting distinct regulatory programs orchestrating the tissue-specific activity of *CG42668* (**Fig. 3h, i**). Furthermore, muscle cells showed additional intra-lineage variation, resulting in a bi-modal distribution of allelic rates (**Fig. 3j**). Despite the presence of strong inter- and intra-lineage variation (Qdiff10 = 0.32), this effect is obscured in a bulk-level analysis (scDALI-Hom *P* = 0.6).

More globally, allele-specific effects are stronger at distal regulatory elements (potential enhancers) compared to promoter-proximal regions, both for heterogeneous (one-sided Mann-Whitney U test, *P* = 5.6×10^−5^) as well as homogeneous (one-sided Mann-Whitney U test, *P* = 2.17×10^−26^) imbalances (**Fig. 3k**). Furthermore, imbalances are significantly more common at distal versus proximal regions (**Fig. 3l**). Similarly to what has been observed in bulk ATAC-seq data sorted by time-matched developmental stages^17^, this difference is only moderate for discoveries by scDALI-Hom (two-sided Binomial test P = 0.02), with about 61% of significant regions being found at proximal regions compared to 62% of all tested peaks. Interestingly, however, we find this effect to be markedly more pronounced for heterogeneously imbalanced regions (two-sided Binomial test *P* = 2.15×10^−10^), with only 47% of peaks discovered by scDALI-Het being located near gene promoters.

To further characterize heterogeneous imbalances, we used scDALI to assess differential lineage effects, testing for differences in mean allelic rates between each lineage and all remaining cells (**Methods**). Briefly, this test can be formulated under the scDALI-Het framework, replacing the continuous cell state kernel with a block-diagonal matrix to indicate lineage membership. Unsurprisingly, the frequency of significant imbalances by lineage (FDR < 0.1) largely resembled the overall read count distribution, suggesting that differences in the number of discoveries across lineages can to a large extent be explained by difference in detection power rather than biological factors (**Fig. 3j, Supp. Fig. 10a**). For the majority of peaks, allele-specific variation is attributable to one or two differentially imbalanced lineages (72%), however, 11% of peaks showing differences between three of four lineages (**Fig. 3k**). Interestingly, for 17% of scDALI-Het discoveries, allele-specific effects do not differentiate any single lineage, indicating the presence of significant intra-lineage variation, for example due to variation in developmental time.

### Identification of sites with heterogeneous allelic imbalance linked to developmental time

Developmental time is a major driver of variation in our dataset and therefore a promising predictor of allele-specific changes within lineages. We applied scDALI to test for time-specific allelic imbalances within muscle, the lineage with the largest number of cells, using the pseudo-temporal ordering estimated by the VAE model as a cell state representation. Leveraging the scDALI framework, we design a kernel capturing both linear and nonlinear (polynomial) temporal dependencies (**Methods**). Out of 363 peaks with significant heterogeneous allelic imbalance that are accessible in muscle (mean total allelic count within lineage < 0.1), scDALI identified 69 (19%) peaks with significant time-specific effects (FDR < 0.1; **Fig. 4a, b**). Notably, 27% of these peaks with time-specific allelic imbalance did not show any lineage-specific effects (**Fig. 4c**). As an example, region chr2R:13675707-13676707 discussed above (**Fig. 3f**) does indeed exhibit strong time-specific imbalances (**Fig. 4e, Fig. 3g**), consistent with the observed intra-lineage variation specifically in muscle cells.

**Figure 4.**
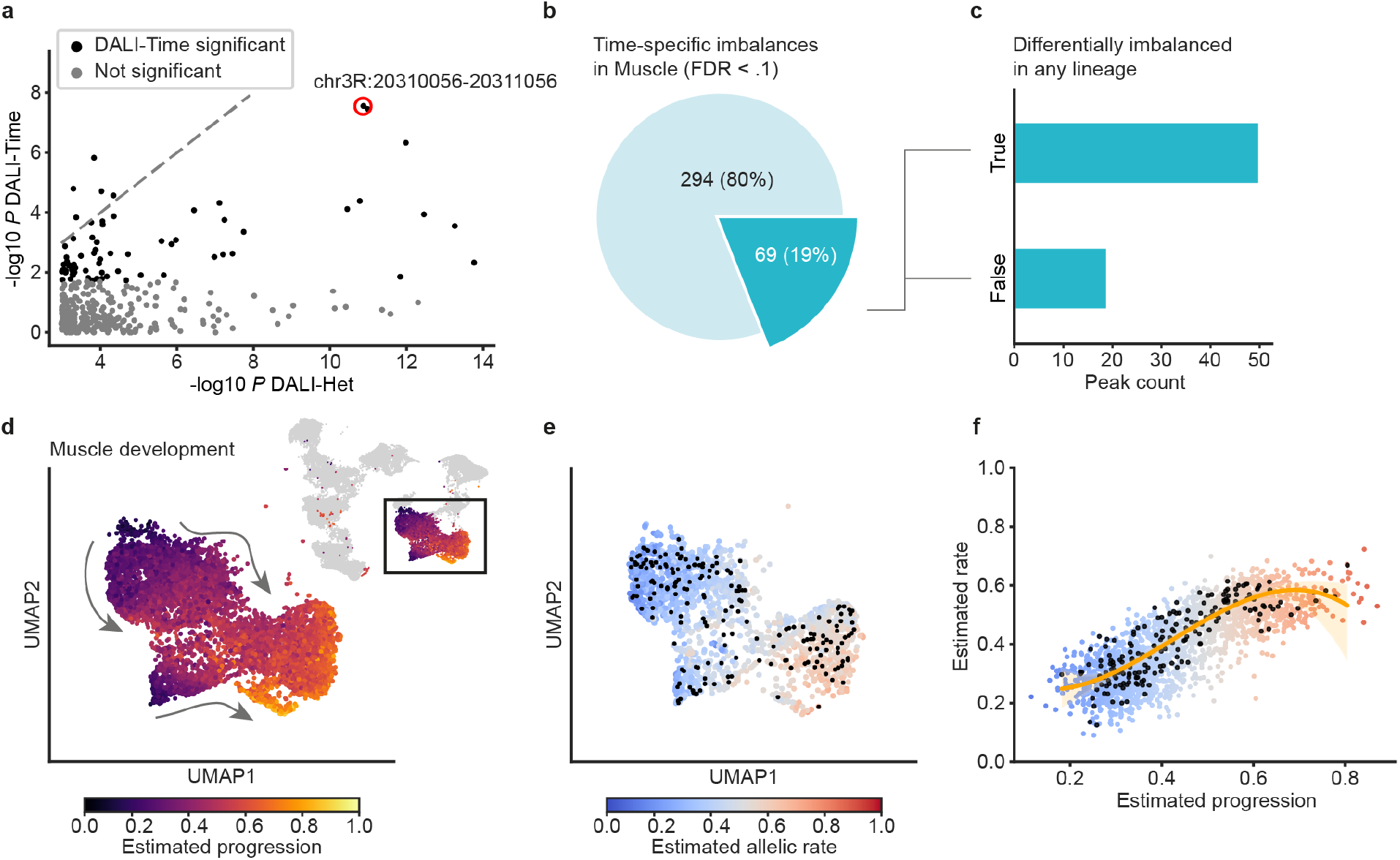
scDALI-Het identifies time-specific intra-lineage variation. **(a)** Scatter plot of negative log P-values for scDALI-Het versus a scDALI test for time-specific variation in the muscle population. Red circle highlights region chr2R:13675707-13676707. **(b)** Of 363 peaks identified by scDALI-Het that are accessible in the muscle, 19% showed significant temporal effects (FDR < 0.1). **(c)** The majority of peaks with a time-specific effect in the muscle did not show significant differential allelic imbalance between lineages. **(d)** Temporal order for the muscle lineage estimated by the variational autoencoder model. **(e, f)** Estimated allelic rates across time for region chr2R:13675707-13676707. Black dots denote cells with observed allele-specific counts in this region.

## Discussion

The majority of disease associated variants impact non-coding regions, disrupting the function of regulatory elements such as enhancers and promoters. As enhancers regulate when and where genes are expressed, genetic variation within enhancers naturally has cell type-specific effects. However, capturing and understanding these genetic effects is an enormous challenge. Resolving these effects to specific cell types using classical Quantitative trait loci (QTL) mapping would require FACS sorting different cell types from a heterogeneous tissue across a large panel of individuals, a huge task that is often impossible as specific markers for cell isolation are not available for many cell types and transitions. Single cell genomics methods can overcome these issues, but there is currently no computational framework to model the functional impact of genetic variation in regulatory regions within and across cell clusters from single cell data.

To address this, we developed scDALI, a computational framework to identify genetic effects from single-cell ATAC sequencing data in an unbiased fashion. Our model provides a principled strategy for exploiting two independent signals that can be obtained from the same sequencing experiment: (1) total accessibility, which we use to derive cell types and states and identify their regulatory elements, and (2) allele-specific quantifications of genetic effects within those regulatory elements. Combining these two measurements allowed us to test for both pervasive, homogeneous imbalance and cell state-specific heterogeneous effects, without the need to define cell types or cell states *a priori*. We applied scDALI to data generated from an F1 cross design, assaying dynamic and discrete changes in allele-specific chromatin accessibility of developing *Drosophila melanogaster* embryos, a naturally very heterogeneous sample. Our model discovers thousands of imbalanced regions, hundreds of which show distinct cell state-specific effects. About half of the regulatory regions with allelic imbalance in specific cell types are not detectable in a pseudo-bulk analysis, as opposing effects cancel out across the cell state space. Although the total number of discoveries with heterogeneous effects is relatively modest, we expect this to increase dramatically as the number of profiled cells by sciATAC increases. Even with the numbers profiled here, our analysis identified genetic effects at a number of characterized developmental enhancers. scDALI can dissect this heterogeneity at different resolutions, by assessing the distribution of allelic activity both between and within lineages or cell types. In particular, we find that developmental time is an important contributor to intra-lineage variation of allelic imbalance, pinpointing developmental stage-specific enhancers. Furthermore, our analysis revealed that allele-specific effects are significantly stronger and more common at distal elements (putative enhancers) compared to promoter-proximal regions. Notably, these differences are markedly more pronounced among peaks with heterogeneous (tissue-specific) imbalances compared to homogeneous effects and previous results on bulk-sequencing data^17^.

Our findings underline that heterogeneous allelic imbalance provides valuable information about regulatory activity beyond the analysis of cell state-specific signals of accessibility. Single-cell technologies not only improve detection power for heterogeneous effects, but also enable us for the first time to map allele-specific effects to cellular states in an unbiased manner.

While our approach uncovers many novel putative regulatory regions, it also has its limitations. Most notably, we do not explicitly include genotyping information which prevents our model from mapping potential causal variants. While in principle, it is possible to combine allelic analyses with genotype data to fine-map causal variants^7–10^, this would require larger numbers of unique genotypes. The required multi-individual single-cell sequencing studies are only beginning to emerge and scDALI could be extended to leverage such variation.

Understanding to what degree allele-specific effects replicate at different molecular layers remains another important direction of future research. Our method is not limited to the application to chromatin accessibility data, but readily generalizes to other count-based assays, including single-cell RNA-seq and single-cell epigenetics data such as DNA methylation. New multi-omics methods can obtain both DNA accessibility and transcriptomic measurements from the same single cell. The integration of these different dimensions of allelic imbalance across both modalities will be an important area for future work, that may help to relate the functional impact of genetic variation in enhancers to their target gene’s expression.

## Methods

A complete methods section, including experimental procedures and the statistical analysis is provided as **Supplementary methods**.

## Data availability

All raw sequencing data have been submitted to the EMBL-EBI ArrayExpress database (https://www.ebi.ac.uk/arrayexpress/) and are available under accession number E-MTAB-10240. Processed data, including the F1 sciATAC peaks per genotype and stage, can all be downloaded from http://furlonglab.embl.de/data.

## Code availability

The scDALI framework including the VAE model for cell state inference is available under an open-source license from https://github.com/PMBio/scdali. Code to reproduce the specific analyses is available from https://github.com/PMBio/scdali_analyses.

## Acknowledgements

The authors thank members of the Furlong and Stegle labs for helpful comments and discussions. T.H. received support from the German Cancer Research Center International PhD Program. This work was technically supported by the EMBL Flow Cytometry, Genomics and Protein Expression and Purification Core Facilities, and financially supported by core funding from EMBL (to E.E.F. and O.S), the German Cancer Research Center (O.S.) and the European Research Council (ERC) grant agreement ID: 787611 (DeCRyPT) to E.E.F and grant agreement ID: 810296 (DECODE) to O.S..

## Author contributions

T.H. developed and implemented scDALI.

T.H. & S.S. analysed data.

T.H. & S.S. generated figures.

S.S. adapted the sciATAC-seq protocol with advice from J.R.

S.S. & J.R. generated sciATAC-seq data.

B.Z. collected the F1 embryos.

T.H., S.S., E.F. & O.S. interpreted results.

T.H. S.S., E.F. & O.S. wrote the manuscript with input from all authors.

E.F. & O.S. supervised the project.

## References

1. Li, X. et al.The impact of rare variation on gene expression across tissues.Nature 550, 239–243 (2017).

2. Ferraro, N. M. et al.Transcriptomic signatures across human tissues identify functional rare genetic variation.Science 369, (2020).

3. GTEx Consortium. The GTEx Consortium atlas of genetic regulatory effects across human tissues.Science 369, 1318–1330 (2020).

4. Cannavò, E. et al.Genetic variants regulating expression levels and isoform diversity during embryogenesis.Nature 541, 402–406 (2017).

5. Cuomo, A. S. E. et al.Single-cell RNA-sequencing of differentiating iPS cells reveals dynamic genetic effects on gene expression.Nat. Commun. 11, 810 (2020).

6. Jerber, J. et al.Population-scale single-cell RNA-seq profiling across dopaminergic neuron differentiation.Nat. Genet. 53, 304–312 (2021).

7. Kumasaka, N., Knights, A.J. & Gaffney, D. J. Fine-mapping cellular QTLs with RASQUAL and ATAC-seq.Nat. Genet. 48, 206–213 (2016).

8. Knowles, D. A. et al.Allele-specific expression reveals interactions between genetic variation and environment.Nat. Methods 14, 699–702 (2017).

9. Sun, W. A statistical framework for eQTL mapping using RNA-seq data.Biometrics 68, 1–11 (2012).

10. van de Geijn, B., McVicker, G., Gilad, Y. & Pritchard, J. K. WASP: allele-specific software for robust molecular quantitative trait locus discovery.Nat. Methods 12, 1061–1063 (2015).

11. Sun, M. & Zhang, J. Allele-specific single-cell RNA sequencing reveals different architectures of intrinsic and extrinsic gene expression noises.Nucleic Acids Res. 48, 533–547 (2020).

12. Jiang, Y., Zhang, N. R. & Li, M. SCALE: modeling allele-specific gene expression by single-cell RNA sequencing. Genome Biol. 18,74(2017).

13. Fan, J., Wang, X., Xiao, R. & Li, M. Detecting cell-type-specific allelic expression imbalance by integrative analysis of bulk and single-cell RNA sequencing data.bioRxiv (2020) 10.1101/2020.08.26.267815.

14. Benaglio, P. et al.Mapping genetic effects on cell type-specific chromatin accessibility and annotating complex trait variants using single nucleus ATAC-seq.bioRxiv (2020) doi:10.1101/2020.12.03.387894.

15. Chen, H. et al.Assessment of computational methods for the analysis of single-cell ATAC-seq data.Genome Biol. 20, 241 (2019).

16. Saelens, W., Cannoodt, R., Todorov, H. & Saeys, Y. A comparison of single-cell trajectory inference methods.Nat. Biotechnol. 37, 547–554 (2019).

17. Floc’hlay, S. et al.Cis-acting variation is common across regulatory layers but is often buffered during embryonic development.Genome Res. 31, 211–224 (2020).

18. Svensson, V., Teichmann, S.A. & Stegle, O. SpatialDE: identification of spatially variable genes.Nat. Methods 15, 343–346 (2018).

19. Buettner, F. et al.Computational analysis of cell-to-cell heterogeneity in single-cell RNA-sequencing data reveals hidden subpopulations of cells.Nat. Biotechnol. 33, 155–160 (2015).

20. Lin, X. Variance component testing in generalised linear models with random effects.Biometrika 84, 309–326 (1997).

21. Zhang,D.& Lin, X. Hypothesis testing in semiparametric additive mixed models.Biostatistics 4, 57–74 (2003).

22. Kingma,D. P. & Welling, M. Auto-Encoding Variational Bayes.arXiv preprint arXiv:1312.6114(2013).

23. Lopez, R., Regier, J., Cole, M.B.,Jordan,M. I. & Yosef, N. Deep generative modeling for single-cell transcriptomics.Nat. Methods 15, 1053–1058 (2018).

24. Xiong, L. et al.SCALE method for single-cell ATAC-seq analysis via latent feature extraction.Nat. Commun. 10, 4576 (2019).

25. Cusanovich, D. A. et al.The cis-regulatory dynamics of embryonic development at single-cell resolution.Nature 555, 538–542 (2018).

26. Traag, V. A.,Waltman,L. & van Eck, N. J. From Louvain to Leiden: guaranteeing well-connected communities.Sci. Rep. 9, 5233 (2019).

27. Wolf, F. A., Angerer,P. & Theis, F. J. SCANPY: large-scale single-cell gene expression data analysis.Genome Biol.19,15 (2018).

28. Reddington, J. P. et al.Lineage-Resolved Enhancer and Promoter Usage during a Time Course of Embryogenesis.Dev. Cell 55, 648–664.e9 (2020).

