## Supplementary Information for "scDALI: Modelling allelic heterogeneity of DNA accessibility in single-cells reveals context-specific genetic regulation"


**Supplementary tables** **2**

**Supplementary figures** **3**

### Supplementary tables

Supplementary table 1 | Summary of scDALI test results provided as a separate data file (supplementary_table1.xlsx). Contained are peaks that were significant at 0.1 FDR for any scDALI test. The fields are as follows:

| | |
|---|---|
| chr | Peak coordinates. |
| start | |
| end | |
| n_cells | Number of cells with nonzero counts. |
| rate | Empirical rate across all cells (sum of reads mapping to the maternal haplotype divided by the total number of reads). |
| qdiff10 | Effect size for heterogeneous allelic imbalance (**Methods**). Only provided for peaks with scdalihet_p_adj < 0.1. |
| scdalihet_p | scDALI-Het p-value. |
| scdalihom_p | scDALI-Hom p-value. |
| scdalijoint_p | scDALI-Joint p-value. |
| scdalihet_p_adj | Benjamini-Hochberg adjusted scDALI-Het p-value. |
| scdalihom_p_adj | Benjamini-Hochberg adjusted scDALI-Hom p-value. |
| scdalijoint_p_adj | Benjamini-Hochberg adjusted scDALI-Joint p-value. |
| scdalijoint_rho | scDALI-Joint $\rho$(estimated relative extent of heterogeneous imbalance, **Methods**). |

### Supplementary figures

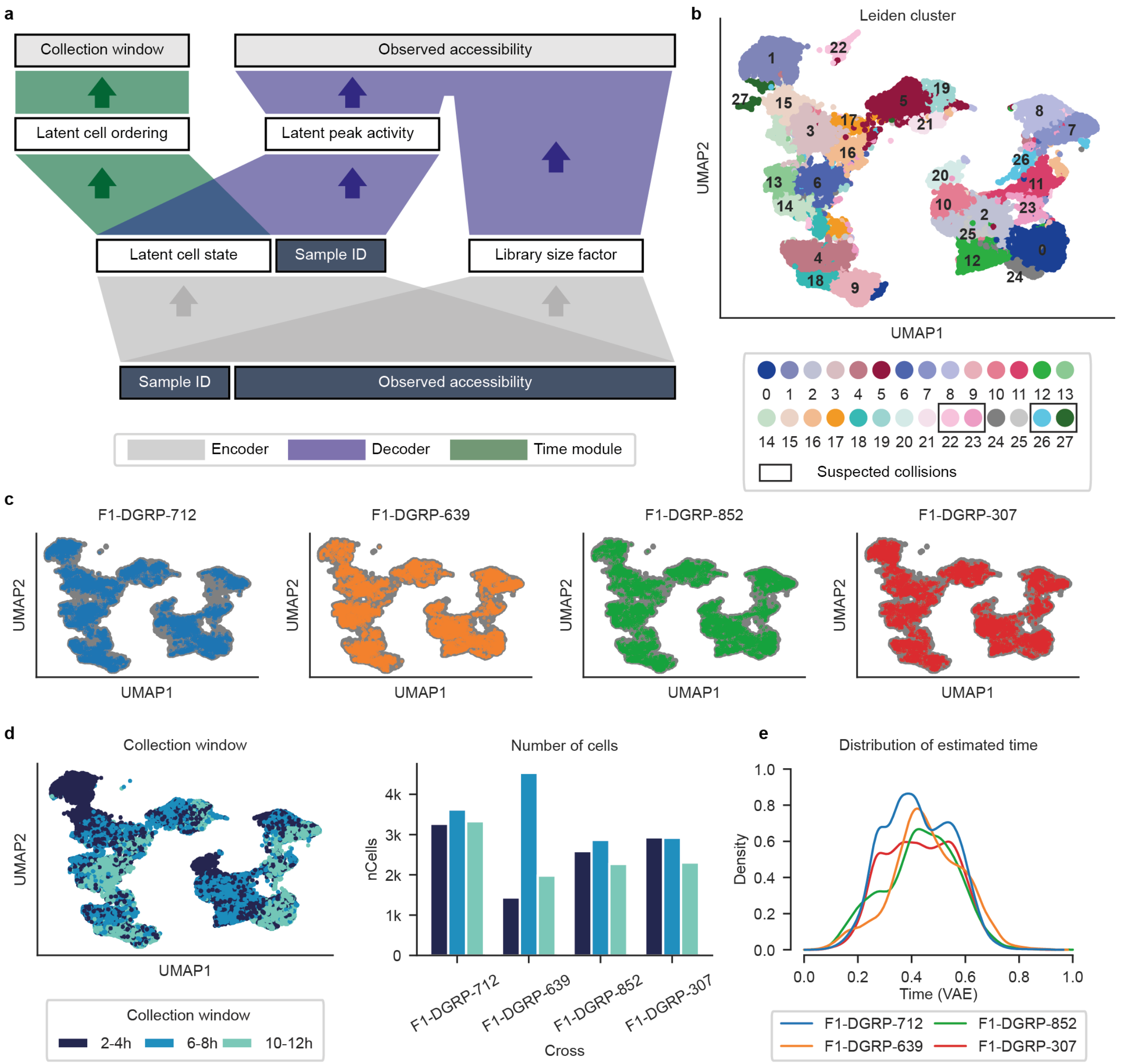


**Supplementary Figure 1 | sci-ATAC Variational autoencoder (VAE). (a)** Model architecture. Colors highlight the three major building blocks. An encoder network maps observed (total) accessibility counts into a lower dimensional cell state space and disentangles library size variation and batch/sample identity. A decoder network first maps cell states and sample ids to a latent peak activity score, indicating the relative “openness” of each peak. Latent peak activity scores are then combined with cell-specific size factors to form a size-factor adjusted Bernoulli likelihood for the observed data. A temporal module infers a continuous pseudo-temporal ordering by mapping cell states to discrete ordered timepoints using an ordinal likelihood model. **(b)** 2-dimensional UMAP visualization of the 8-dimensional VAE cell state representation colored by Leiden clusters. Clusters 22, 23, 26 and 27 were suspected to correspond to barcode collisions and later discarded (**Methods**). **(c)** UMAP visualization as in **b** with individual panels highlighting the cells from each cross. The VAE uniformly integrates all four F1 crosses in a joint cell state space. **(d)** Left: UMAP embedding colored by the (observed) time point of embryo collection, revealing a coarse-grained temporal trajectory. Right: Number of cells associated with each embryo collection window after filtering cells with very high (more than 99% quantile) or very low counts (less than 10% quantile) in each timepoint. **(e)** Distribution of estimated continuous time for individual cells as inferred by the VAE model for each cross.

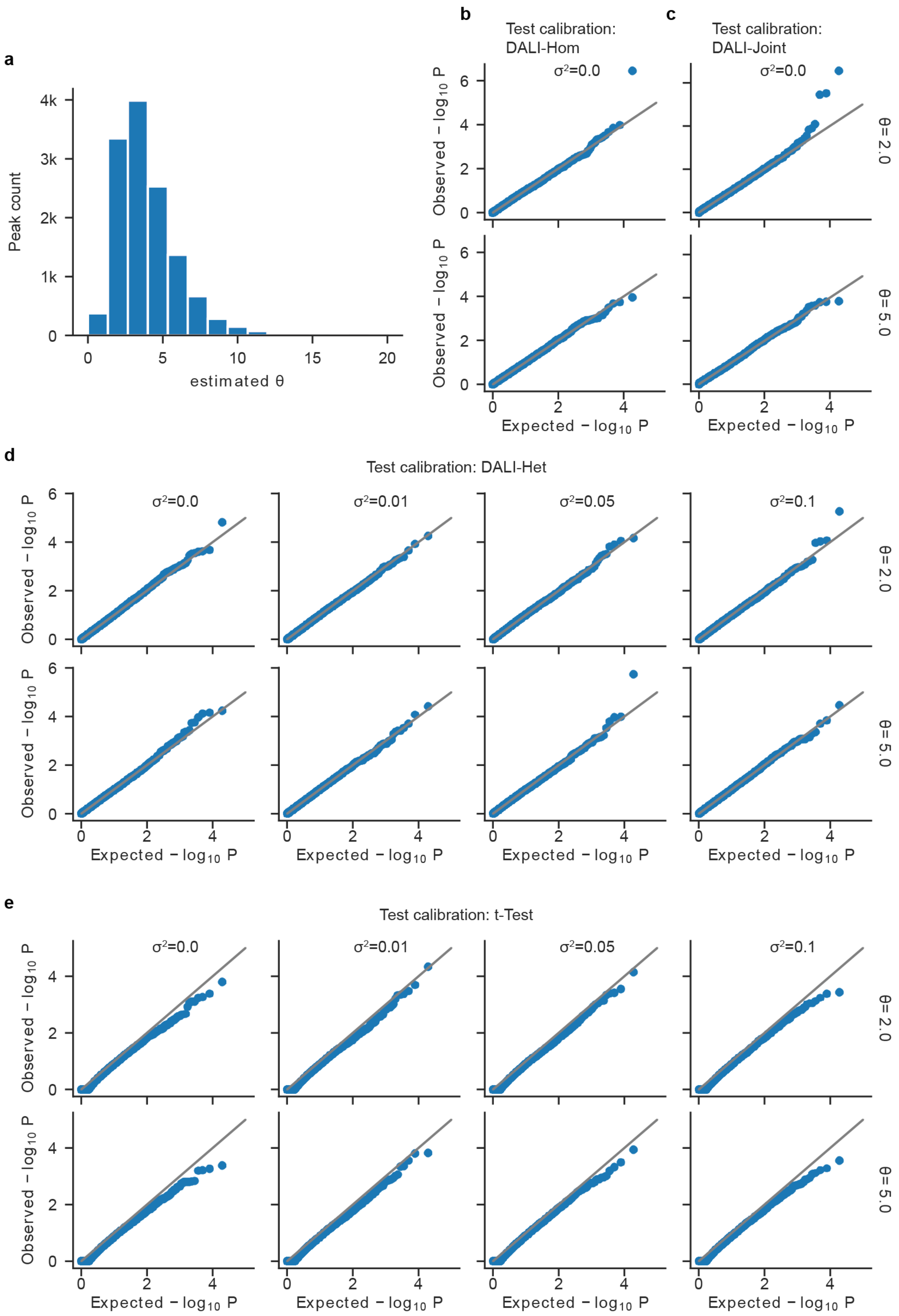


**Supplementary Figure 2 | Test calibration on simulated data. (a)** Histogram of estimated overdispersion parameters for individual peaks of chromatin accessibility for F1 cross

F1-DGRP-712 (10,220 cells and 12,861 peak), using a Beta-Binomial model without any cell state-specific information. **(b-e)** QQ plots to assess empirical calibration of alternative tests on data simulated from a Beta-Binomial model, with constant mean sampled from a Normal distribution with mean zero and variance $\sigma^2$(columns) and two alternative levels of overdispersion $\theta$ (rows). A nonzero value of $\sigma^2$ introduces homogeneous but no heterogeneous imbalance. Shown are results from 12,861 tests, where total counts correspond to observed data as in **a**. scDALI-Joint and scDALI-Het models were provided with a cell state kernel obtained from a VAE model trained on real data (**Supp. Fig. 3, Methods**). Considered were **(b)** scDALI-Hom, **(c)** scDALI-Joint, **(d)** scDALI-Het and **(e)** a one-vs-all t-test testing for differences in mean allelic rates between Leiden clusters inferred from real data.

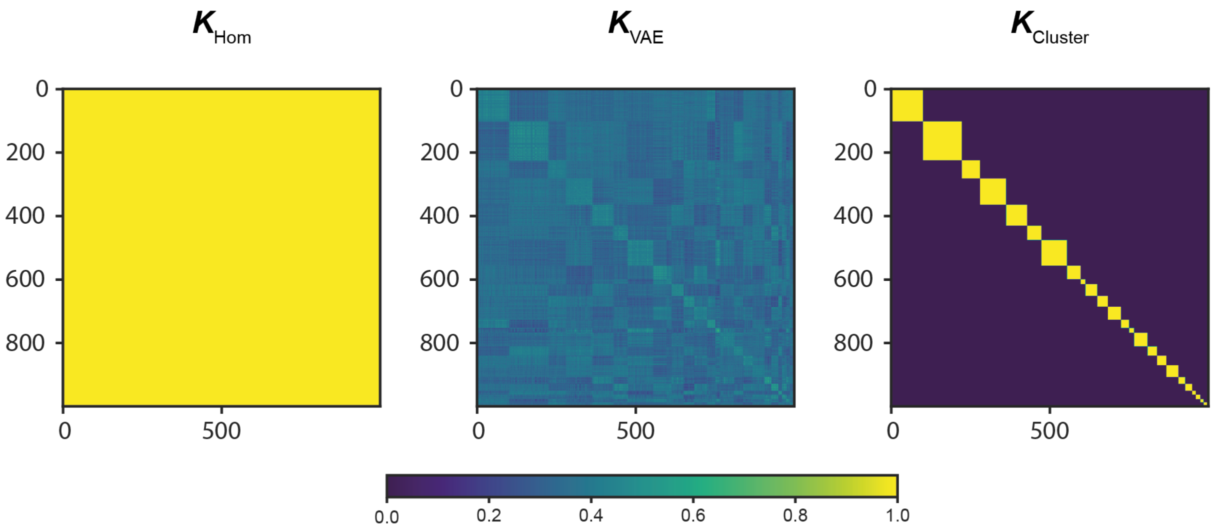


**Supplementary Figure 3 | Kernels for the simulation procedure.** Kernel matrices for simulation experiments. Left: Homogeneous kernel (matrix of ones). Middle: Cell state kernel based on the inner product similarity in the VAE cell state space for 10,220 cells from cross F1-DGRP-712 (**Methods, Supp. Fig. 1a**). Right: Discrete cluster kernel; a block diagonal matrix indicating Leiden cluster membership (N=25 clusters, **Supp. Fig. 1b**). All kernels are subsampled to 1,000 cells for visualization purposes.

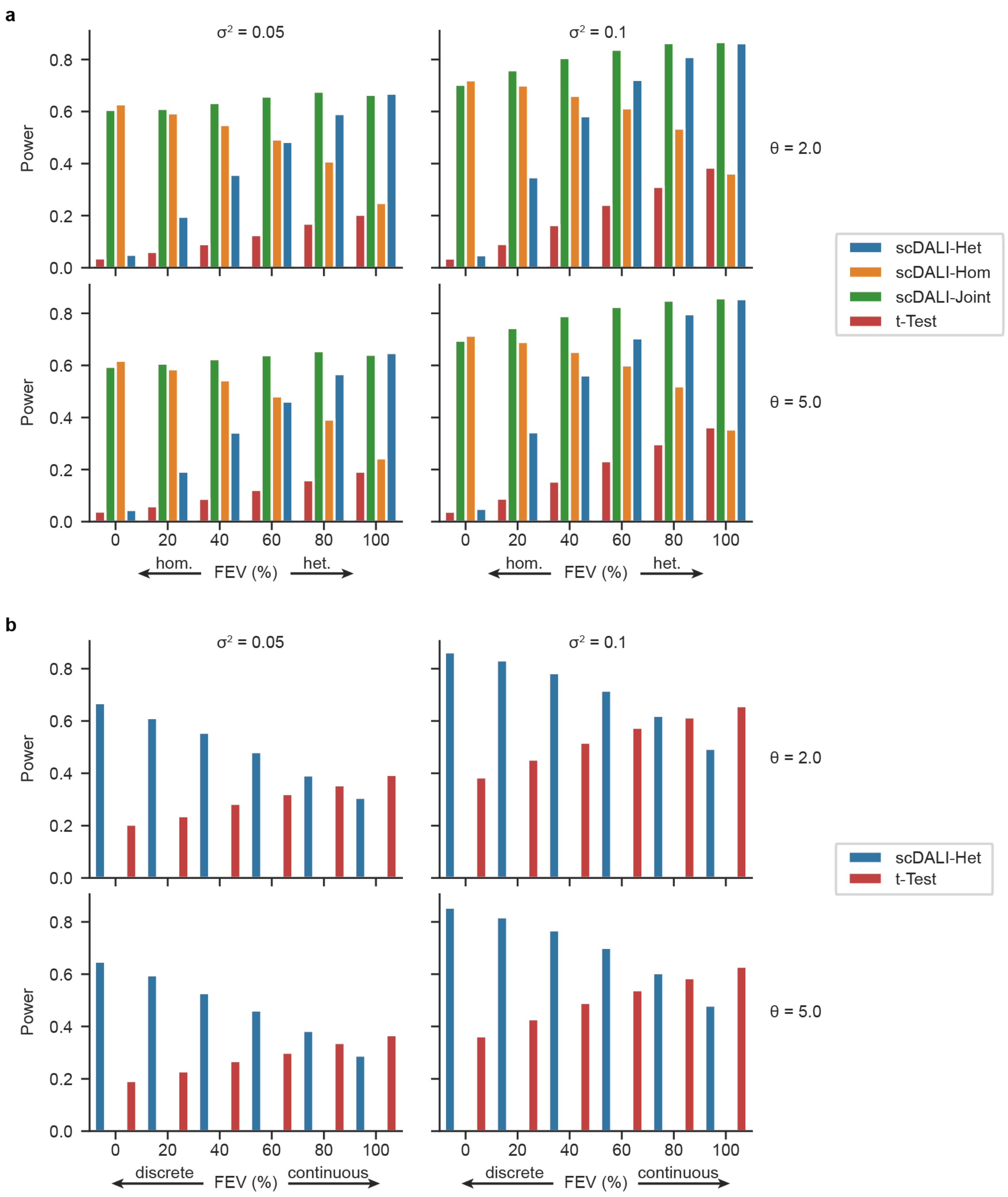


**Supplementary Figure 4 | Additional results from the power assessment on simulated data.** Alternative counts were generated from the scDALI model, when considering different levels of variance explained by the simulation kernel ($\sigma^2$, columns) for two levels of overdispersion ($\theta$, rows). Shown are results as in **Fig. 1b-e**, when **(a)** varying the extent of simulated heterogeneous vs. homogeneous allelic imbalance and **(b)** varying between discrete and continuous cell states (fraction of explained variance, FEV, **Methods**). Barplots show the fraction of peaks discovered by scDALI-Het, scDALI-Hom and scDALI-Joint.

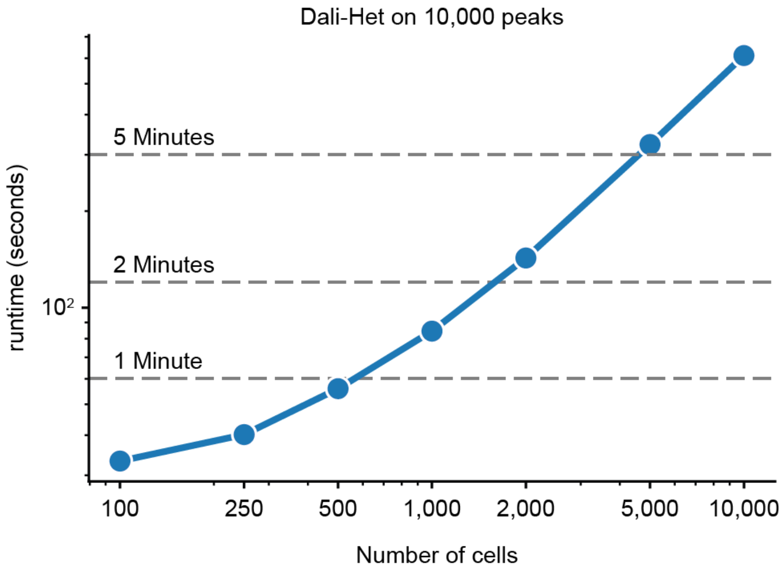


**Supplementary Figure 5 | DALI-Het runtime analysis.** Empirical runtimes for 10,000 tests of randomly sampled peaks for cross F1-DGRP-712 for increasing number of cells.DALI-Het scales linearly with the number of cells. Cells with non-zero total allelic counts were sampled with replacement to ensure the same number for each peak. Runtimes were evaluated on a 2018 MacBook Pro with 2,3 GHz Quad-Core Intel Core i5 processor.

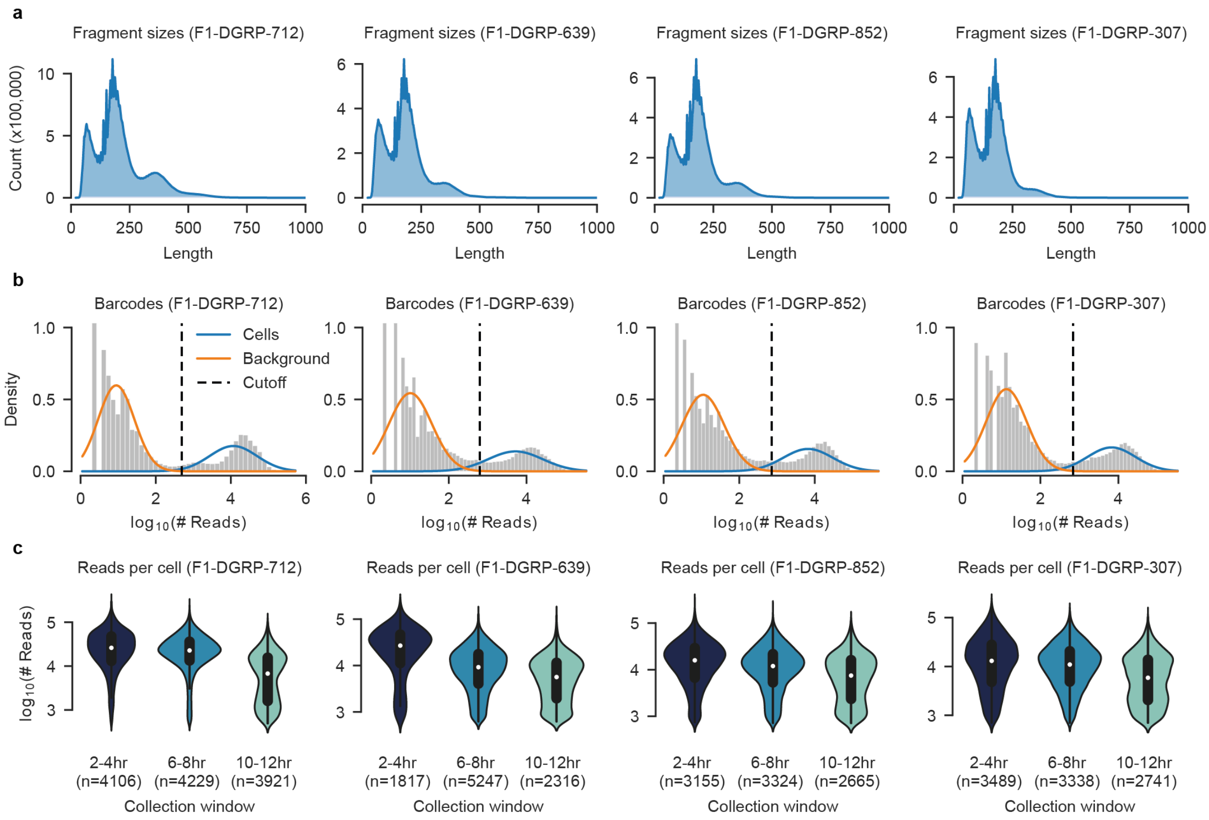


**Supplementary Figure 6 | QC for *Drosophila melanogaster* sci-ATAC-seq data. (a)** Fragment size distribution for all four crosses. The multi-modal distribution corresponds to the expected nucleosome banding pattern. **(b)** Distribution of log-total read counts per barcode. A two-component Gaussian mixture model is fitted to discern background signal from genuine cells (orange and blue lines). The dotted line represents the cutoff corresponding to the 95% posterior probability of cells belonging to the foreground mixture component. **(c)** Violin plots showing the read count distribution for cells identified in **b** stratified by the embryo collection window (2-4, 6-8 and 10-12 hours after egg laying).

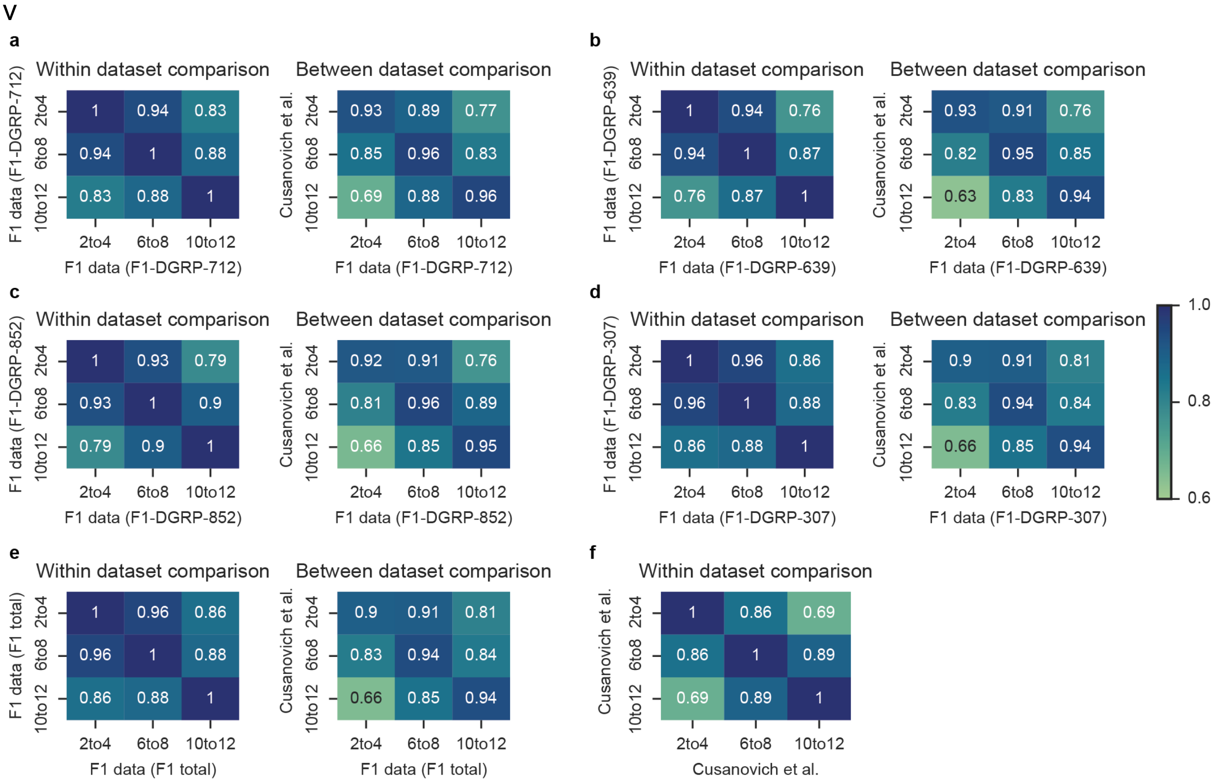


**Supplementary Figure 7 | Comparison with published sci-ATAC data. (a-d)** Pearson correlation of pseudo-bulk accessibility profiles across ATAC peaks, both within our dataset as well as compared to published, time-matched sci-ATAC data (Cusanovich et al. 2018) for each cross. Matching timepoints are highly correlated between both datasets. **(e)** Pearson correlation for the combined dataset comprising all four crosses. **(f)** Within dataset comparison for the Cusanovich et al. data.

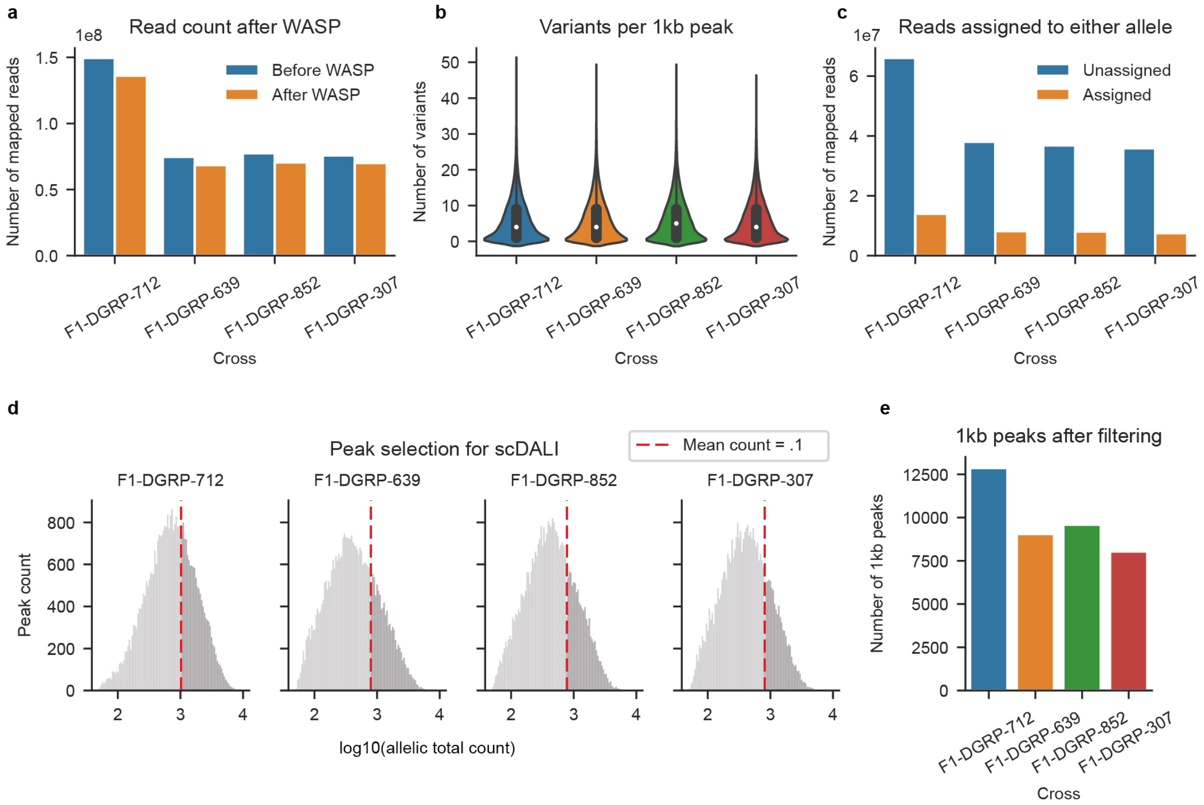


**Supplementary Figure 8 | QC for allele-specific quantifications. (a)** Read counts before and after applying the WASP pipeline to reduce reference mapping biases. **(b)** Number of variants across 1kb windows centered on peaks in each cross. **(c)** Number of assigned and unassigned reads for each cross. About 20% of all reads overlapping 1kb windows centered on peaks can be assigned to either allele. **(d)** Peak selection for the scDALI tests. Shown are histograms of the allelic total read counts (total number of reads that can be assigned to either allele). The dotted red line indicates a mean allelic total count per 1kb peak of .1, which was used as a filtering cutoff. **(e)** Number of 1kb peaks per cross after filtering. The combined set of 39,530 peaks was tested for allelic imbalance using scDALI.

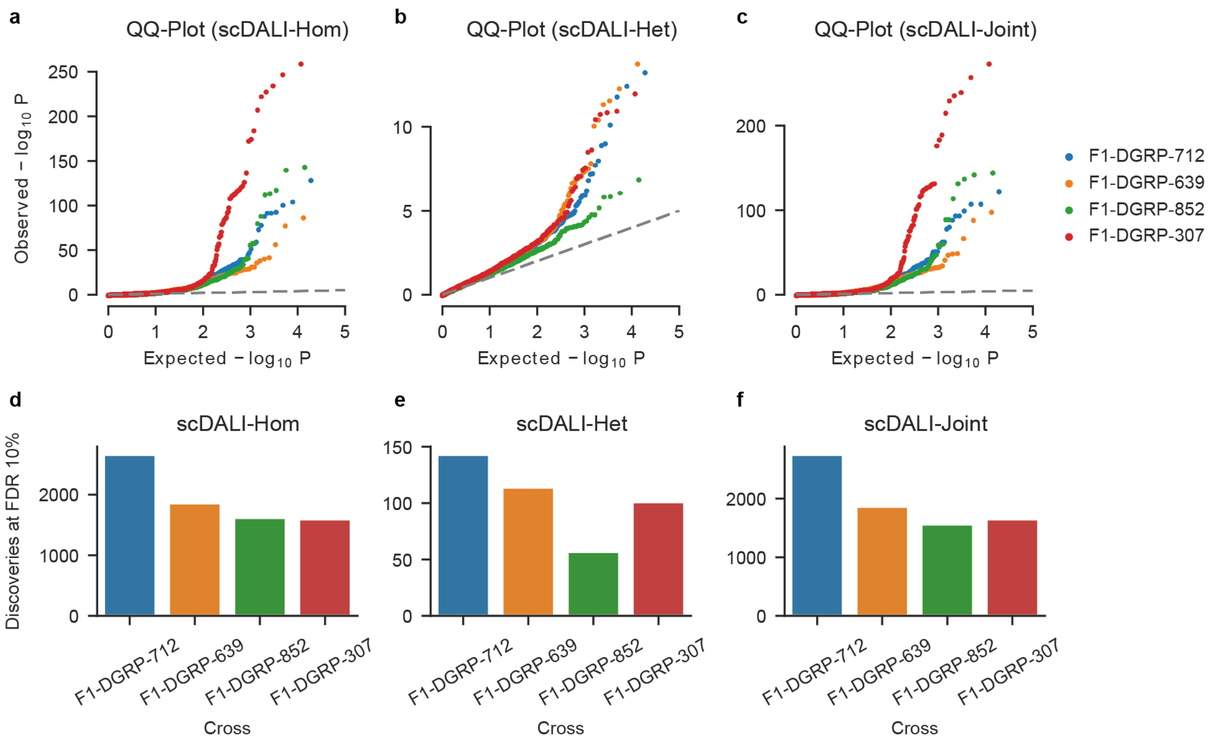


**Supplementary Figure 9 | DALI discoveries. (a-c)** QQ Plots for scDALI-Hom, scDALI-Het and scDALI-Joint applied to allele-specific ATAC-seq of developing Drosophila embryos. Colors indicate four different crosses. **(d-f**) Number of discoveries for each test stratified by cross.

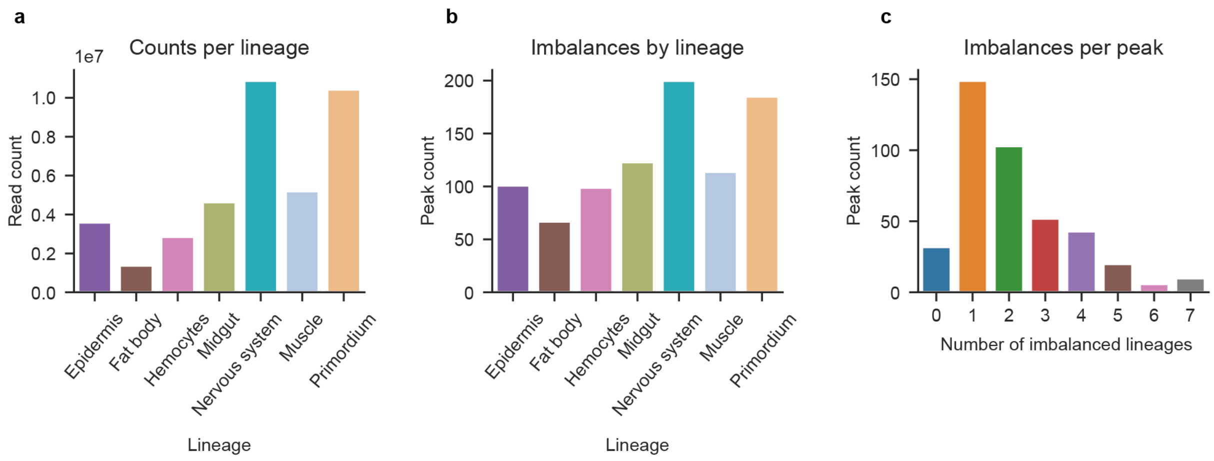


**Supplementary Figure 10 | Additional results from the by lineage analysis.** Considered were peaks with significant heterogeneous imbalances (adjusted scDALI-Het P < 0.1) **(a)** Total read count per lineage. **(b-c)** scDALI-Hom applied to discover (homogeneous) allelic imbalances per lineage. **(b)** Number of peaks showing allelic imbalances (FDR < 0.1). The number of discoveries strongly resembles the total read count per lineage in **a.** **(c)** Number of imbalances per peak. The majority of peaks with heterogeneous allele-specific effects show imbalances in one or two lineages.
