## Supplementary Methods for "scDALI: Modelling allelic heterogeneity of DNA accessibility in single-cells reveals context-specific genetic regulation"

T. Heinen, S. Secchia, J. P. Reddington, B. Zhao,
E. E. M. Furlong, O. Stegle

**5 Software availability** **17**

### 1 The scDALI model

scDALI extends the frequently used Beta-Binomial observation model for allele-specific counts in bulk-sequencing data [1, 2, 3, 4], by accounting for cell state-specific effects.

We model the relationship between cell states and allelic rates for a given genomic region using a hierarchical Bayesian model. For cells $i = 1, \ldots, n$, let $a_i$ be the number of reads mapping to the the maternal haplotype and $d_i$ be the total number of reads (for ease of notation, we omit the dependence on the region of interest). We will initially assume that the measurements for each cell correspond to independent trials, and introduce the basic Beta-Binomial model. In section 1.2 we then discuss how the scDALI framework can be used to capture cell state specific covariances. While our terminology will focus on chromatin accessibility data, the same modeling principles generalize to other count-based single-cell assays including single-cell RNA-seq.

#### 1.1 A Beta-Binomial model for allelic imbalance

Assume we are studying the allele-specific accessibility of a particular region among cells from a homogeneous population. That is, all cells share the same underlying allelic rate $p$, the average accessibility of the maternal haplotype. Then, given the observed total number of reads $d_i$, we can regard $a_i$ as a draw from a binomial distribution

$$a_i \,|\, d_i \sim \mathrm{Bin}(d_i, p). \tag{1}$$

Real data, however, shows more variability than is to be expected under a binomial model. This is because cellular populations are rarely entirely homogeneous (e.g. due to cell cycle effects) and allele-specific counts are affected by additional technical and biological sources of variation. One can account for this by making $p$ itself a random variable. A common choice is the Beta distribution [1, 2, 3, 4]

$$p \sim \mathrm{Beta}(\theta^{-1}\mu, \theta^{-1}(1-\mu)), \tag{2}$$

leading to a compound distribution with closed-form density. Here, $\mu$ denotes the mean allelic rate and $\theta$ is an overdispersion parameter modulating the amount of extra-binomial variance. For small $\theta$, the Beta-Binomial distribution approaches a Binomial distribution. Conversely, when $\theta$ is large, draws will resemble a Bernoulli distribution with counts coming almost exclusively from either allele.

##### 1.1.1 Parameter estimation from single-cell data

While the Beta-Binomial distribution does not allow for closed-form maximum-likelihood estimates, efficient numerical optimization algorithms are available [5]. Estimating parameters from bulk sequencing data is challenging, as the number of replicates is often limited. To lower the estimation uncertainty, further assumptions are usually required, such as a shared mean-variance relationship between genomic regions [1]. Single-cell sequencing assays, on the other hand, provide large numbers of cells / samples and in principle allow for direct estimation of $\theta$ and $\mu$ separately for each region. Note that we are still assuming that the allelic rates in different cells are independent and identically distributed (i.i.d.). This assertion, however, is unlikely to hold in practice as molecular traits are correlated across cell types and states. In the next section, we therefore extend eq. (1-2) to account for cellular heterogeneity.

#### 1.2 Capturing heterogeneous imbalance

Building on Gaussian process (GP) regression [6], we capture cell state-specific allelic variation by introducting a latent $n$-dimensional Gaussian variable

$$\mathbf{u} \sim \mathcal{N}(\mathbf{1} \cdot \alpha, \sigma_{het}^2 \mathbf{K}). \tag{3}$$

Here, $\mathbf{u}$ denotes the vector of allelic rates on the logit scale, which is coupled to the Beta-Binomial observation model for allelic counts introduced in section 1.1 as follows

$$\mu_i = g^{-1}(u_i) \tag{4}$$

$$a_i \,|\, \mu_i, d_i \sim \text{Beta-Binomial}(\theta^{-1}\mu_i, \theta^{-1}(1 - \mu_i)), \tag{5}$$

where $g$ is the logit link function, $g(x) = \log(\frac{x}{1-x})$. The fixed effect $\alpha$ models homogeneous imbalance (here $\alpha = 0$ corresponds to an allelic rate of 1/2), while cell state covariances are encoded in a kernel matrix $\mathbf{K} \in \mathbb{R}^{n \times n}$. The scaling parameter $\sigma_{het}^2$ determines the total variance explained by cell state-specific effects. In particular, $\sigma_{het}^2 = 0$ implies $\mu_i = \mu = g^{-1}(\alpha)$ and we recover the basic Beta-Binomial model (1-2).

Throughout this paper we use a linear kernel function $\mathbf{K}(\mathbf{E}) = \mathbf{E}\mathbf{E}^T$ where $\mathbf{E} \in \mathbb{R}^{n \times k}$ is a matrix of $k$-dimensional cell states based on the total (non-allele-specific) chromatin accessibility profiles for each cell. For example, $\mathbf{E}$ could contain the coordinates for each cell in a $k$-dimensional embedding of the total accessibility matrix or a one-hot encoding for $k$ cell clusters. In section 3 we describe how we infer cell states across different individuals or batches using a variational autoencoder model. Our choice of a linear kernel function is primarily motivated by computational efficiency (see section 1.3.1). Note, however, that we may still account for non-linear effects by transforming or combining cell state dimensions (see section 4.6.2 for an example). Alternatively, more general associations can be captured using non-linear kernel functions [6].

##### 1.3 Statistical significance testing

The scDALI model specified in eq. (3-5) uses two parameters to capture allelic imbalances: The fixed effect $\alpha$ encodes *homogeneous* imbalance affecting all cells equally across the state space, while the variance component $\sigma^2_{het}$ determines the magnitude of cell state-specific, i.e. *heterogeneous* effects. We consider three different null and alternative hypothesis, for which efficient score-based tests are derived in the following sections:

**scDALI-Hom** Presence of *homogeneous* allelic imbalance (section 1.3.3)

$$H_0^{hom} : \alpha = 0 \text{ vs. } H_1^{hom} : \alpha \neq 0. \tag{6}$$

**scDALI-Het** Presence of *heterogeneous* allelic imbalance (section 1.3.1)

$$H_0^{het} : \sigma^2_{het} = 0 \text{ vs. } H_1^{het} : \sigma^2_{het} > 0. \tag{7}$$

**scDALI-Joint** Presence of either *homogeneous* or *heterogeneous* allelic imbalance (section 1.3.2)

$$H_0^{joint} : \alpha = 0 \text{ and } \sigma^2_{het} = 0 \text{ vs. } H_1^{joint} : \alpha \neq 0 \text{ or } \sigma^2_{het} > 0. \tag{8}$$

###### 1.3.1 scDALI-Het

To test for heterogeneous allelic imbalance, we need to assess whether $\sigma^2_{het} > 0$. One possibility is to use a likelihood ratio test (LRT) [7]. However, the LRT requires to fit parameters for both the null and alternative models which is computationally expensive. Furthermore, the null hypothesis places $\sigma^2_{het}$ on the boundary of the parameter space, and will result in a likelihood-ratio test statistic that does not follow a chi-square distribution asymptotically. Instead, we implement an efficient score-based testing procedure [8, 9, 10, 11], scaling linearly with the number of cells if the cell state kernel $\mathbf{K}$ is of low rank.

The derivation of the score statistic for the model defined in Eq. (4-5) largely follows [9] and [10]. Consider the maternal allelic ratios $r_i = a_i/d_i$ under the model defined in (3-5). Conditioned on the vector of random effects $\mathbf{u}$, the $r_i$ are independently following a Beta-Binomial distribution with

$$\mathbb{E}[r_i \,|\, u_i] = \mu_i \tag{9}$$

$$\text{Var}(r_i \,|\, u_i, d_i) = V(\mu_i, d_i) = \frac{1}{d_i}\mu_i(1-\mu_i)\frac{\theta^{-1} + d_i}{\theta^{-1} + 1}. \tag{10}$$

Let $(\hat{\alpha}_0, \hat{\theta}_0)$ be a maximum-likelihood estimate of $(\alpha, \theta)$ under the null model $H_0^{het}$ and let $\hat{\mu}_0 = g^{-1}(\hat{\alpha}_0)$.

Similar to [9], we define a null covariance matrix for the Beta-Binomial distribution

$$\mathbf{W}_0 = \text{diag}((V(\hat{\mu}_0, d_i)g'(\hat{\mu}_0)^2)^{-1}) = \text{diag}((\frac{1}{d_i\hat{\mu}_0(1-\hat{\mu}_0)}\frac{\hat{\theta}_0^{-1}+d_i}{\hat{\theta}_0^{-1}+1})^{-1}) \tag{11}$$

$$= \text{diag}(d_i\hat{\mu}_0(1-\hat{\mu}_0)\frac{\hat{\theta}_0+1}{d_i\hat{\theta}_0+1}), \tag{12}$$

and the projection matrix accounting for the fixed effect

$$\mathbf{P}_0 = \mathbf{W}_0 - \mathbf{W}_0\mathbf{1}(\mathbf{1}^T\mathbf{W}_0\mathbf{1})^{-1}\mathbf{1}^T\mathbf{W}_0. \tag{13}$$

Following [9], we define the score-based statistic

$$Q = \frac{1}{2}\tilde{\mathbf{r}}_0^T\mathbf{P}_0^T\mathbf{K}\mathbf{P}_0\tilde{\mathbf{r}}_0 \tag{14}$$

where $\tilde{\mathbf{r}}_0$ is the working vector with entries

$$(\tilde{r}_0)_i = \mathbf{1}\cdot\hat{\alpha}_0 + g'(\hat{\mu}_0)(r_i - \hat{\mu}_0). \tag{15}$$

One can show that the distribution of $Q$ under the null model $H_0^{het}$ can be approximated by a weighted sum of independent $\chi_1^2$ random variables [12]

$$\sum_k \lambda_k\chi_{1,k}^2, \tag{16}$$

where $\lambda_k$ are the ordered non-zero eigenvalues of $\frac{1}{2}\mathbf{P}_0^{T/2}\mathbf{K}\mathbf{P}_0^{1/2}$. While computing the eigenvalues of a general $n \times n$ matrix has a computational complexity of $O(n^3)$, a linear kernel $\mathbf{K} = \mathbf{E}\mathbf{E}^T$ typically allows for a much more efficient implementation. Note that for all matrices $\mathbf{A}$, it holds that eigenvalues$(\mathbf{A}^T\mathbf{A}) =$ eigenvalues$(\mathbf{A}\mathbf{A}^T)$ [13]. Therefore, we have

$$\text{eigenvalues}(\frac{1}{2}\mathbf{P}_0^{T/2}\mathbf{K}\mathbf{P}_0^{1/2}) = \text{eigenvalues}(\frac{1}{2}(\mathbf{E}^T\mathbf{P}_0^{1/2})^T(\mathbf{E}^T\mathbf{P}_0^{1/2})) \tag{17}$$

$$= \text{eigenvalues}(\frac{1}{2}(\mathbf{E}^T\mathbf{P}_0^{1/2})(\mathbf{E}^T\mathbf{P}_0^{1/2})^T) \tag{18}$$

$$= \text{eigenvalues}(\frac{1}{2}\mathbf{E}^T\mathbf{P}_0\mathbf{E}), \tag{19}$$

which can be computed in $O(k^3)$, where the cell state dimensionality $k$ is typically much smaller than the number of cells $n$. To evaluate the distribution function and obtain p-values we use Davies method [14, 15] as implemented in `limix`[1].

[1] `https://github.com/limix/chiscore`

##### 1.3.2 scDALI-Joint

To implement a joint test capable of identifying either heterogeneous or homogeneous allelic imbalance, we adapt the approach originally developed for SKAT-O [10] and more recently extended in [11]. For a given null maternal rate $\mu_0$, let $\alpha_0 = g^{-1}(\mu_0)$. We model homogeneous imbalance $\alpha$ as a second random effect $\alpha \sim \mathcal{N}(\alpha_0, \sigma^2_{hom})$ such that (3) becomes

$$\mathbf{u} \sim \mathcal{N}(\mathbf{1} \cdot \alpha_0, \sigma^2_{hom}\mathbf{1}\mathbf{1}^T + \sigma^2_{het}\mathbf{K}). \tag{20}$$

Equivalently, one can write

$$\mathbf{u} \sim \mathcal{N}(\mathbf{1} \cdot \alpha_0, \sigma^2_{tot}[(1-\rho)\mathbf{1}\mathbf{1}^T + \rho\mathbf{K}]) \tag{21}$$

where $\sigma^2_{tot} = \sigma^2_{het} + \sigma^2_{hom}$ denotes the total variance explained by allele-specific effects, while $\rho = \sigma^2_{het}/\sigma^2_{tot} \in [0,1]$ corresponds to relative extent of heterogeneous imbalance (**Fig. 2d**). This allows us to reformulate scDALI-Joint as a one-parameter variance component test

$$H_0^{joint} : \sigma^2_{tot} = 0 \text{ vs. } H_1^{joint} : \sigma^2_{tot} > 0 \text{ (scDALI-Joint)}. \tag{22}$$

For given $\rho$, we can test this hypothesis using the same score-based framework described above for scDALI-Het, assuming a modified kernel matrix $\mathbf{K}_\rho = [(1-\rho)\mathbf{1}\mathbf{1}^T + \rho\mathbf{K}]$ and fixing $\alpha = \alpha_0$ in (3). As $\rho$ is unknown in practice, we perform a grid search and combine the resulting p-values:

1. Compute p-values $p_{\rho_r}$ for $\rho_r$ from a pre-defined grid of values in $[0,1]$. As in [10, 11], we replace the exact Davies method with the modified moment matching approximation [15, 16] to improve computational efficiency when evaluating the null distribution (16).

2. Determine the test statistic $T = \min_r p_{\rho_r}$ and compute the final p-value. Details on the exact form and estimation of the associated null distribution can be found in [11].

##### 1.3.3 scDALI-Hom

As a special case, we can use the scDALI framework to test for homogeneous allelic imbalance by fixing $\rho = 0$ in eq. (21) (corresponding to no cell state-specific effects). Alternatively, one can employ a likelihood ratio test

$$LLR = 2 \cdot \log \frac{p(\mathbf{a} \,|\, \mathbf{d}, \mu = \hat{\mu}_1, \theta = \hat{\theta}_1)}{p(\mathbf{a} \,|\, \mathbf{d}, \mu = \mu_0, \theta = \hat{\theta}_0)}, \tag{23}$$

where $p(\cdot)$ denotes the likelihood function for the model described in (1-2), $\mu_0$ is the null allelic rate and all other parameters are estimating using maximum-likelihood. P-values can be calculated using the fact that $LLR$ asymptotically follows a chi-square distribution with one degree of freedom [17].

#### 1.4 Allelic rate estimation and effect size computation

To estimate the landscape of allele-specific chromatin accessibility we approximate the scDALI model (eq. (3-5)) using Gaussian process regression [6]. Specifically, (using the notation of section 1.2) we model the empirical rates $r_i = a_i/d_i$ as

$$r_i \sim \mathcal{N}(\mathbf{1} \cdot \alpha, \sigma_{het}^2 \mathbf{K} + \eta^2 \mathbf{I}), \tag{24}$$

where $\eta$ captures residual Gaussian noise independent of the cell state component. Equivalently, we can write $r_i = u_i + \epsilon$ where $\epsilon \sim \mathcal{N}(0, \eta)$ and $\mathbf{u}$ is defined in eq. (3). The hyper-parameters $\alpha, \sigma_{het}^2$ and $\eta^2$ along with posterior approximations are fitted using sparse variational inference [18] as implemented in GPflow [19].

While the estimate for $\alpha$ provides a measure of pervasive, homogeneous allelic imbalance, we can use the posterior mean of the latent variable $\mathbf{u}$ as an estimate for the cell-specific allelic rate. In particular, we define a measure of effect size or statistical dispersion for heterogeneous effects

$$\text{Qdiff10} = Q_{0.9} - Q_{0.1} \tag{25}$$

where $Q_{0.9}$ and $Q_{0.1}$ denote the 0.9 and 0.1 quantiles of the estimated posterior mean for $\mathbf{u}$ (**Fig. 3c**).

#### 2 Evaluation of scDALI on simulated data

We simulated allele-specific counts from the scDALI model formalized in eq. (21) using observed allelic total counts (10220 cells and 12861 peaks) and inferred cell state representations (section 4.4) from real sci-ATAC-seq data of developing *Drosophila melanogaster* embryos (cross F1-DGRP-712, see section 4). Specifically, we considered a linear kernel $\mathbf{K}_{vae}$ based on the 8-dimensional cell states inferred by the VAE model (section 3) and a block diagonal matrix $\mathbf{K}_{cluster}$ indicating Leiden cluster membership (section 4.4, **Supp. Fig. 3**). Both the scDALI-Het and scDALI-Joint tests were performed using the VAE kernel $\mathbf{K}_{vae}$. To asses the degree of overdispersion present in the data, we initially fitted a basic Beta-Binomial model to the observed allele-specific counts using no additional cell state information (eq. (1-2, **Supp. Fig. 2a**)). Based on the histogram of estimated values, we ran all simulations at two different levels of overdispersion ($\theta \in \{2, 5\}$).

We assessed the statistical calibration of all three scDALI tests by simulating from a model assuming no heterogeneous imbalance and different levels of pervasive (cell state independent) effects ($\rho = 0$, $\sigma^2_{tot} \in \{0, 0.01, 0.05, 0.1\}$; **Supp. Fig. 1b-d**). As an additional baseline, we considered a one-vs-all t-test to detect heterogeneous allelic imbalance in the form of differences between the mean allelic rates of each Leiden cluster and the remaining cells (minimum p-value across all clusters, Bonferroni corrected for the number of clusters; **Supp. Fig. 1e**).

We compared power to detect homogeneous vs. heterogeneous effects by simulating data varying the relative extent of heterogeneous imbalance ($\rho \in \{0, 0.2, 0.4, 0.6, 0.8, 1\}$), as well as total variance explained by allele-specific effects ($\sigma^2_{tot} \in \{0.01, 0.05, 0.1\}$, **Fig. 1c, Supp. Fig. 4a**). Significance was determined at an $\alpha$-level of 0.05. Lastly, we assessed power to detect discrete vs. contiuous heterogeneous effects, using a weighted combination of the VAE kernel and the Leiden cluster kernel

$$\mathbf{K} = \alpha \mathbf{K}_{cluster} + (1 - \alpha)\mathbf{K}_{vae}. \tag{26}$$

In this scenario we assumed no additional homogeneous effects ($\rho = 1$) and again considered a range of weights and kernel scaling parameters. ($\alpha \in \{0, 0.2, 0.4, 0.6, 0.8, 1\}$, $\sigma^2_{tot} \in \{0.1, 0.05, 0.1\}$, **Fig. 1d, Supp. Fig. 4b**).

### 3 The cell state variational autoencoder

To test for heterogeneous allelic imbalance, we require a suitable representation of cell types or states. Here, we implement a variational autoencoder model (VAE) [20, 21] to infer a lower-dimensional embedding of the total (non-accessibility) profiles. Variations of VAE models have been widely applied to model single-cell transcriptome meassurements [22, 23, 24, 25, 26] and more recently been extended to model chromatin accessibility data [26]. Our model is most closely related to scVI [22], a VAE capable of integrating scRNA-seq data across different individuals or batches while accounting for library-size variation. Different from scVI, however, we use a likelihood model tailored to the near-binary nature of single-cell ATAC-seq data. Furthermore, we integrate sampling times for developmental datasets to estimate a continuous pseudo-temporal ordering from few available timepoints. Our model can be decomposed into three sub-modules (**Supp. Fig. 1a**): The decoder network (section 3.1), representing the generative process for observed accessibility profiles, the temporal classifier (section 3.3) and the encoder network for inferring the posterior distribution over latent variables (section 3.2).

#### 3.1 A generative model for single-cell chromatin accessibility

Let $\mathbf{x}_i \in \{0, 1\}^m$ be the binarized accessibility vector for $m$ peaks in cell $i$ and $c_i$ be the batch / individual identity. The probabilistic generative model underlying the decoder module can be formalized as follows:

$$\mathbf{z}_i \sim \mathcal{N}(\mathbf{0}, \mathbf{I}) \tag{27}$$

$$l_i \mid c_i \sim \text{LogNormal}(\mu_l(c_i), \sigma_l^2(c_i)) \tag{28}$$

$$\boldsymbol{\rho_i} = f_\rho(\mathbf{z}_i, c_i) \tag{29}$$

$$x_{ij} \mid \rho_{ij}, l_i \sim \text{Bernoulli}(1 - (1 - \rho_{ij})^{l_i}) \tag{30}$$

Here, $\mathbf{z}_i \in \mathbb{R}^k$, are the latent, low-dimensional cell state representations and $l_i \in \mathbb{R}_{>0}$ is a cell-specific size-factor variable capturing variation in sequencing depth [22]. The prior parameters $\mu_l(c_i)$ and $\sigma_l^2(c_i)$ are chosen to be maximum-likelihood estimates based on the total number of reads per cell in each cross. cell states along with observed batch ids are mapped to $m$-dimensional peak activities $\boldsymbol{\rho}_i \in [0, 1]^m$, $\sum_j \rho_{ij} = 1$, representing the relative "openness" of each peak in cell $i$. The mapping is realized by a neural network $f_\rho$ with trainable parameters. By providing both $f_\rho$ and the encoder network (see section 3.2 below) with $c_i$, the model is encouraged to disentangle batch-specific effects and cell state representations [22, 27] (**Supp. Fig. 1c**). The full distribution over observed accessibility profiles is obtained by applying the scaling factor to the peak activities. If $l_i$ were the true (discrete) number of reads per cell, $1 - (1 - \rho_{ij})^{l_i}$ would correspond to the probability of observing at least one read in peak d. However, to simplify the inference process, we place a continuous log-normal prior on $l_i$.

#### 3.2 Incorporating temporal information

For the data considered in this paper, coarse temporal information in the form of embryo collection windows is available (**Supp. Fig. 1d**). Here, we integrate these time stamps with the observed ATAC-seq data to inform the latent-space inference, correct time measurement errors and learn a continuous ordering of cells from few available labels. More specifically, assume the time label $y_i$ for cell $i$ takes on one of $t$ ordered values. If $t$ is small, it is difficult to accurately estimate the temporal scale at which cell state changes take place. Instead, we model the relative order of cells as a function of the cell state $\mathbf{z}_i$, using an ordinal likelihood model. Formally, we assume $y_i \in \{1, 2, \ldots, t\}$ and define [28]

$$p(y_i \,|\, \mathbf{z}_i) = \Phi(w_{y_i} - f_y(\mathbf{z}_i)) - \Phi(w_{y_i - 1} - f_y(\mathbf{z}_i)) \tag{31}$$

where $\Phi$ denotes the cumulative distribution function of the standard normal distribution, $w_0 = -\infty$, $w_t = \infty$ and $w_1, \ldots, w_{T-1}$ are trainable parameters such that $w_i < w_{i+1}$. The function $f_y$ maps cell states to pseudo-temporal values along the real axis and is chosen to be a simple linear model. Intuitively, $p(y_i \,|\, \mathbf{z}_i)$ corresponds to the probability of sampling a value from the interval $(w_{y_i - 1}, w_{y_i})$ under a Normal distribution with mean $f_y(\mathbf{z}_i)$ and unit variance. By allowing for Gaussian noise around the latent time $f_y(\mathbf{z}_i)$, we can account for measurement error in the labeling process. Note that the $w_i$ form a contiguous segmentation of the real line which enforces ordinal constraints.

Guided by both observed time stamps and cell state proximities, our model infers a high resolution pseudo-temporal trajectory $f_y$, allowing us to order cells according to their developmental progression. We optimize all parameters jointly with the other modules, thereby incentivizing our model to learn a cell state representation that is informed by and supports the observed time labels (**Fig. 2b, Supp. Fig. 1e**).

#### 3.3 Variational inference and encoder networks

Even though the posterior distribution over latent variables $\mathbf{z}_i, l_i$ is intractable, we can approximate it using variational inference. That is, we define parametric families of distributions $q(l_i \,|\, \mathbf{x}_i, c_i)$ and $q(\mathbf{z}_i, \,|\, \mathbf{x}_i, c_i)$ and maximize the following variational lower bound on the marginal likelihood

$$\begin{aligned} \log p(x) \geq\ & E_{q(\mathbf{z}, \,|\, \mathbf{x}, c) q(l \,|\, \mathbf{x}, c)}[\log p(\mathbf{x} \,|\, \mathbf{z}, l, c) p(y \,|\, \mathbf{z})] \\ & - \mathrm{KL}[q(\mathbf{z} \,|\, \mathbf{x}, c) \,||\, p(\mathbf{z})] \\ & - \mathrm{KL}[p(l \,|\, \mathbf{x}, c) \,||\, p(l)]. \end{aligned} \tag{32}$$

Here $\mathrm{KL}(\cdot \,||\, \cdot)$ denotes the KL divergence and we have made the simplifying assumption that $l_i \,|\, \mathbf{x}_i, c_i$ and $\mathbf{z}_i, \,|\, \mathbf{x}_i, c_i$ are independent (mean-field variational inference). A key idea of the variational autoencoder framework is the use of neural networks known as *recognition networks* or *encoders* to map observed data to the parameters of the variational distributions [20]. Following [22], we use a Gaussian posterior for $\mathbf{z}_i$ and a Log-Normal posterior for $l_i$, allowing for closed-form solutions for the KL terms. Furthermore, we can make use of a specific sampling scheme known as the reparameterization trick [20], facilitating training both prior and variational parameters in an end-to-end fashion.

##### 3.4 Practical considerations

We implement all neural networks using batch normalization [29] and ReLU activation functions between hidden layers. We use ADAM [30] for iterative stochastic optimization and approximate the expectation in eq. (32) using 5 Monte-Carlo samples from the posterior distribution. To avoid over-regularization at the early stages of training, we apply a scaling factor to the KL term to modulate its influence. More specifically, we found that shrinking the KL term in epoch $i$ by a factor of $\frac{\mathrm{i}}{25}$ worked well on our data. Hyper-parameter for the sci-ATAC-seq data of developing *Drosophila melanogaster* embryos were tuned by maximizing the held-out log likelihood on 20% of the cells. The final choices are shown in table 1.

Table 1: VAE hyper-parameters

| Parameter | Value |
|---|---|
| Cell state dimension $k$ | 8 |
| Hidden layers for the encoder $q(\mathbf{z}_i \mid \mathbf{x}_i, c_i)$ | [256, 128] |
| Hidden layers for the encoder $q(l_i, \mid \mathbf{x}_i, c_i)$ | [256] |
| Hidden layers for the decoder $f_\rho(\mathbf{z}_i, c_i)$ | [64, 128] |
| Hidden layers for time module $f_z(\mathbf{z}_i)$ | None |

### 4 Analysis of allelic imbalance in Drosophila Melanogaster

We generated *Drosophila melanogaster* F1 hybrids by crossing females from a common maternal virgnizier line with males from four different inbred lines from the Drosophila melanogaster genetic reference panel (DGRP) [31, 1]. Embryos were collected in two hours windows (2-4 hours, 6-8 hours and 10-12 hours after egg laying) as described in [1].

#### 4.1 Generation of Tn5 transposomes for combinatorial indexing

Hyperactive Tn5 transposase was purified by the EMBL Protein Expression and Purification facility as previously described [32] and stored at $-20°$C in storage buffer (25 mM Tris pH 7.5, 800 mM NaCl, 0.1 mM EDTA, 1 mM DTT, 50% glycerol) until use. Uniquely indexed oligonucleotides from [33] were annealed to common pMENTs oligos (95°C 5 min, cooling to 65°C (0.1°C/sec), 65°C 5 min, cooling to 4°C (0.1°C/sec)) to generate indexed transposons that were then loaded onto purified Tn5 by incubation at 23°C with constant shaking at 350 rpm for 30 minutes. The loaded Tn5 transposomes were diluted 1:10 (final 0.02 mg/ml) in nuclease-free water and used immediately for tagmentation.

#### 4.2 Generation and sequencing of sci-ATAC-seq libraries

Embryo dissociation and nuclear isolation were performed as described previously [33]. Nuclei were flash frozen in liquid nitrogen and stored at $-80°$C until use. Generation of sci-ATAC-seq libraries was performed largely as previously described [33], with minor modifications. The tagmentation reaction was performed by adding two microliters of each of the 96 custom and uniquely indexed Tn5 transposomes and by incubating at 55°C for one hour. After reverse-crosslinking, 5 $\mu$L of forward and reverse indexed primers (from [33]), 7.5 $\mu$L KAPA HiFi DNA Polymerase ReadyMix (Roche) and 0.25 $\mu$L Bst3.0 (NEB) were added to each well. Tagmented DNA was then PCR amplified with the following cycling conditions: 72°C 5 min, 98°C 30 s; 98°C 10 s, 63°C 30 s, 19–22 cycles; 72°C 1 min, hold at 10°C. The optimal number of cycles for each library was determined beforehand by monitoring amplification on a qPCR machine for a set of test wells. Libraries were sequenced on an Illumina NextSeq 500 sequencer High Capacity 150 PE kit as previously described [33].

#### 4.3 Processing of sequencing data

Raw sequencing data was processed based on the pipeline developed in [33]. BCL files were converted to fastq files using bcl2fastq v.2.16 (Illumina). To correct for sequencing or PCR amplification errors, read barcodes were matched again all possible barcodes. In case of an approximate match (Levenshtein distance $< 3$ and distance to next best match $> 2$) the corresponding barcodes were fixed to their presumptive match; all other barcodes were classified as ambiguous or unknown. Barcode correction was followed by adapter trimming [34] and read alignment to the dm6 reference genome using bowtie [35] (with options `-X 2000 -3 1`). After the removal of PCR duplicates, we classified barcodes corresponding to genuine cells from the background by fitting a two-component Gaussian mixture model to the log-transformed read counts per barcode. A cutoff for cell barcodes was determined by requiring that the posterior probability of belonging to the higher read-depth mixing component was greater than 0.95 (**Supp. Fig. 6b, c**).

#### 4.4 Count preprocessing and cell state definition

Chromatin accessibility was quantified in a set of 53,133 peaks of accessibility previously identified from a time-matched sci-ATAC-seq dataset [33] and lifted to the dm6 reference genome[2]. We compared the Pearson correlation between pseudo-bulk aggregates for each collection window both within our dataset as well as between our data and the published reference (**Supp. Fig. 7**). Next, we restricted our analysis to autosomes in order to remove sex-specific biases [33]. For each cross and collection window, we determined the 10% and 99% quantiles of the cell-count distribution and only kept cells whose counts were within those limits, resulting in 35,485 cells in total. Finally, we selected the 25,000 top most accessibile peaks for further analysis. We trained the cell state variational autoencoder on the whole dataset for 30 epochs, using the network architecture specified in table 1.

We used the Scanpy implementation of the Leiden algorithm [36, 37] with a resolution of 1.2 and identified 28 cell clusters in the joint VAE latent space. For each cluster, we computed differentially accessible peaks using logistic regression by predicting cluster labels from the peak activity profiles obtained from the VAE model (see eq. (30)) [38, 37]. We then performed an enrichment analysis for known tissue specific enhancer elements (CAD4 database, [33]) and genes (tissue-specific expression of the nearest gene based on *in situ* hybridization data from the Berekely *Drosophila* Genome Project and FlyBase gene expression annotations) using a Fisher's exact test. Based on these enrichments, each cluster was assigned to one of seven major lineages (**Fig. 2b**). Four clusters with a total of 1,432 cells could not be annotated unambigously and were removed from further analysis (**Fig 1b**), resulting in a final set of 35,485 cells (**Supp. Fig. 1d**).

#### 4.5 WASP pipeline and processing of allele-specific counts

We used existing genotyping information [1] to create cross-specific VCF files, filtering for genetic variants that segregate parental strains. To eliminate reference mapping biases, we applied the WASP pipeline [3], filtering out between 7-8% of mapped reads (**Supp. Fig. 8a**). For each of the 35,485 cells we then

[2]https://github.com/FlyBase/bulkfile-scripts

quantified allele-specific accessibility in 1kb windows centered on each of the 53,133 peaks to overcome the inherent sparsity of the data. Reads aligned within these windows were assigned to either allele, requiring that each read overlapped at least one heterozygous single-nucleotide variant. As it can be challenging to accurately estimate the allelic base rate for sex chromosomes (that is, the overall proportion of female embryos), we excluded these from our analysis. Finally, windows in each cross were filtered by requiring that the mean allelic total count (that is, the sum of reads that could be assigned to either allele) across cells was no smaller than 0.1 (**Supp. Fig. 8d**). This resulted in between 8,040 and 12,861 peaks per cross and a combined set of 39,530 peaks to be tested for allelic imbalance (**Supp. Fig. 8e**).

#### 4.6 scDALI analysis

We applied scDALI to all of the 39,530 peaks to test for heterogeneous (scDALI-Het), homogeneous (scDALI-Hom) and either kind of allelic imbalance (scDALI-Joint). Both scDALI-Het and scDALI-Joint used the 8-dimensional VAE latent space embedding as a cell state representation and a linear kernel function. P-values from each test were Benjamini-Hochberg adjusted to control the false discovery rate (FDR) [39]. For each of the 415 sites with evidence for cell state-specific allelic imbalance (scDALI-Het $P < 0.1$ FDR) we estimated allelic rates for all cells using GP regression (section 1.4). Depending on the number of covered cells for each peak, all models were trained with a maximum of 1,000 inducing points.

##### 4.6.1 By-lineage analysis

To identify variable lineages for each of the 415 peaks with heterogeneous imbalance, we used scDALI-Het with a lineage-specific cell state kernel. Specifically, for each lineage $\mathcal{L}$, we defined the indicator vector $\mathbf{E}_{\mathcal{L}} \in \{0,1\}^n$

$$(\mathbf{E}_{\mathcal{L}})_i = \begin{cases} 1 & \text{cell } i \text{ in lineage } \mathcal{L} \\ 0 & \text{otherwise} \end{cases}. \tag{33}$$

Using a linear kernel $K(\mathbf{E}_{\mathcal{L}}) = \mathbf{E}_{\mathcal{L}}\mathbf{E}_{\mathcal{L}}^T$, then allowed us to test for differences in the mean allelic rate of lineage $\mathcal{L}$ compared to the mean of all other cells. The combined P-values for all lineages were adjusted for multiple testing using the Benjamini-Hochberg correction [39].

##### 4.6.2 Temporal analysis

Using the temporal ordering inferred by VAE model, we employed scDALI-Het to test for changes in allelic imbalance across development in the muscle lineage. To capture non-linear effects, we used a polynomial basis transform. Specifically, let $t_i \in [0,1], i \in \mathcal{L}_{Muscle}$ denote the estimated time for each cell in the muscle lineage. Then we defined

$$\mathbf{E}_{Time} = \begin{bmatrix} \vdots & \vdots & \vdots \\ t_i & t_i^2 & t_i^3 \\ \vdots & \vdots & \vdots \end{bmatrix} \tag{34}$$

and $\mathbf{K}(\mathbf{E}_{Time}) = \mathbf{E}_{Time}\mathbf{E}_{Time}^T$. P-values were adjusted using the Benjamini-Hochberg correction [39].

### 5 Software availability

scDALI is available under an open source license at `https://github.com/PMBio/scDALI`. Code to reproduce the specific analyses presented can be accessed under `https://github.com/PMBio/scDALI_analyses`.

### References

[1] Swann Floc'hlay, Emily S Wong, Bingqing Zhao, Rebecca R Viales, Morgane Thomas-Chollier, Denis Thieffry, David A Garfield, and Eileen EM Furlong. Cis-acting variation is common across regulatory layers but is often buffered during embryonic development. *Genome research*, 31(2):211–224, 2021.

[2] Natsuhiko Kumasaka, Andrew J Knights, and Daniel J Gaffney. Fine-mapping cellular qtls with rasqual and atac-seq. *Nature genetics*, 48(2):206–213, 2016.

[3] Bryce Van De Geijn, Graham McVicker, Yoav Gilad, and Jonathan K Pritchard. Wasp: allele-specific software for robust molecular quantitative trait locus discovery. *Nature methods*, 12(11):1061–1063, 2015.

[4] Wei Sun. A statistical framework for eqtl mapping using rna-seq data. *Biometrics*, 68(1):1–11, 2012.

[5] Thomas Minka. Estimating a dirichlet distribution, 2000.

[6] Christopher KI Williams and Carl Edward Rasmussen. *Gaussian processes for machine learning*, volume 2. MIT press Cambridge, MA, 2006.

[7] Valentine Svensson, Sarah A Teichmann, and Oliver Stegle. Spatialde: identification of spatially variable genes. *Nature methods*, 15(5):343–346, 2018.

[8] Xihong Lin. Variance component testing in generalised linear models with random effects. *Biometrika*, 84(2):309–326, 1997.

[9] Daowen Zhang and Xihong Lin. Hypothesis testing in semiparametric additive mixed models. *Biostatistics*, 4(1):57–74, 2003.

[10] Seunggeun Lee, Mary J Emond, Michael J Bamshad, Kathleen C Barnes, Mark J Rieder, Deborah A Nickerson, ESP Lung Project Team, David C Christiani, Mark M Wurfel, Xihong Lin, et al. Optimal unified approach for rare-variant association testing with application to small-sample case-control whole-exome sequencing studies. *The American Journal of Human Genetics*, 91(2):224–237, 2012.

[11] Rachel Moore, Francesco Paolo Casale, Marc Jan Bonder, Danilo Horta, Lude Franke, Inês Barroso, and Oliver Stegle. A linear mixed-model approach to study multivariate gene–environment interactions. *Nature genetics*, 51(1):180–186, 2019.

[12] Christoph Lippert, Jing Xiang, Danilo Horta, Christian Widmer, Carl Kadie, David Heckerman, and Jennifer Listgarten. Greater power and computational efficiency for kernel-based association testing of sets of genetic variants. *Bioinformatics*, 30(22):3206–3214, 2014.

[13] Kaare Brandt Petersen and Michael Syskind Pedersen. The matrix cookbook.

[14] Robert B Davies. Algorithm as 155: The distribution of a linear combination of $\chi$ 2 random variables. *Journal of the Royal Statistical Society. Series C (Applied Statistics)*, 29(3):323–333, 1980.

[15] Pierre Duchesne and Pierre Lafaye De Micheaux. Computing the distribution of quadratic forms: Further comparisons between the liu–tang–zhang approximation and exact methods. *Computational Statistics & Data Analysis*, 54(4):858–862, 2010.

[16] Huan Liu, Yongqiang Tang, and Hao Helen Zhang. A new chi-square approximation to the distribution of non-negative definite quadratic forms in non-central normal variables. *Computational Statistics & Data Analysis*, 53(4):853–856, 2009.

[17] Samuel S Wilks. The large-sample distribution of the likelihood ratio for testing composite hypotheses. *The annals of mathematical statistics*, 9(1):60–62, 1938.

[18] Michalis Titsias. Variational learning of inducing variables in sparse gaussian processes. In *Artificial intelligence and statistics*, pages 567–574. PMLR, 2009.

[19] Alexander G de G Matthews, Mark Van Der Wilk, Tom Nickson, Keisuke Fujii, Alexis Boukouvalas, Pablo León-Villagrá, Zoubin Ghahramani, and James Hensman. Gpflow: A gaussian process library using tensorflow. *J. Mach. Learn. Res.*, 18(40):1–6, 2017.

[20] Diederik P Kingma and Max Welling. Auto-encoding variational bayes. *arXiv preprint arXiv:1312.6114*, 2013.

[21] Carl Doersch. Tutorial on variational autoencoders. *arXiv preprint arXiv:1606.05908*, 2016.

[22] Romain Lopez, Jeffrey Regier, Michael B Cole, Michael I Jordan, and Nir Yosef. Deep generative modeling for single-cell transcriptomics. *Nature methods*, 15(12):1053–1058, 2018.

[23] Christopher Heje Grønbech, Maximillian Fornitz Vording, Pascal N Timshel, Casper Kaae Sønderby, Tune H Pers, and Ole Winther. scvae: Variational auto-encoders for single-cell gene expression data. *Bioinformatics*, 36(16):4415–4422, 2020.

[24] Valentine Svensson, Adam Gayoso, Nir Yosef, and Lior Pachter. Interpretable factor models of single-cell rna-seq via variational autoencoders. *Bioinformatics*, 36(11):3418–3421, 2020.

[25] Dongfang Wang and Jin Gu. Vasc: dimension reduction and visualization of single-cell rna-seq data by deep variational autoencoder. *Genomics, proteomics & bioinformatics*, 16(5):320–331, 2018.

[26] Lei Xiong, Kui Xu, Kang Tian, Yanqiu Shao, Lei Tang, Ge Gao, Michael Zhang, Tao Jiang, and Qiangfeng Cliff Zhang. Scale method for single-cell atac-seq analysis via latent feature extraction. *Nature communications*, 10(1):1–10, 2019.

[27] Chenling Xu, Romain Lopez, Edouard Mehlman, Jeffrey Regier, Michael I Jordan, and Nir Yosef. Probabilistic harmonization and annotation of single-cell transcriptomics data with deep generative models. *Molecular systems biology*, 17(1):e9620, 2021.

[28] Wei Chu, Zoubin Ghahramani, and Christopher KI Williams. Gaussian processes for ordinal regression. *Journal of machine learning research*, 6(7), 2005.

[29] Sergey Ioffe and Christian Szegedy. Batch normalization: Accelerating deep network training by reducing internal covariate shift. In *International conference on machine learning*, pages 448–456. PMLR, 2015.

[30] Diederik P Kingma and Jimmy Ba. Adam: A method for stochastic optimization. *arXiv preprint arXiv:1412.6980*, 2014.

[31] Trudy FC Mackay, Stephen Richards, Eric A Stone, Antonio Barbadilla, Julien F Ayroles, Dianhui Zhu, Sònia Casillas, Yi Han, Michael M Magwire, Julie M Cridland, et al. The drosophila melanogaster genetic reference panel. *Nature*, 482(7384):173–178, 2012.

[32] Matthew J Rossi, William KM Lai, and B Franklin Pugh. Simplified chip-exo assays. *Nature communications*, 9(1):1–13, 2018.

[33] Darren A Cusanovich, James P Reddington, David A Garfield, Riza M Daza, Delasa Aghamirzaie, Raquel Marco-Ferreres, Hannah A Pliner, Lena Christiansen, Xiaojie Qiu, Frank J Steemers, et al. The cis-regulatory dynamics of embryonic development at single-cell resolution. *Nature*, 555(7697):538–542, 2018.

[34] Anthony M Bolger, Marc Lohse, and Bjoern Usadel. Trimmomatic: a flexible trimmer for illumina sequence data. *Bioinformatics*, 30(15):2114–2120, 2014.

[35] Ben Langmead and Steven L Salzberg. Fast gapped-read alignment with bowtie 2. *Nature methods*, 9(4):357, 2012.

[36] Vincent A Traag, Ludo Waltman, and Nees Jan Van Eck. From louvain to leiden: guaranteeing well-connected communities. *Scientific reports*, 9(1):1–12, 2019.

[37] F Alexander Wolf, Philipp Angerer, and Fabian J Theis. Scanpy: large-scale single-cell gene expression data analysis. *Genome biology*, 19(1):1–5, 2018.

[38] Vasilis Ntranos, Lynn Yi, Páll Melsted, and Lior Pachter. A discriminative learning approach to differential expression analysis for single-cell rna-seq. *Nature methods*, 16(2):163–166, 2019.

[39] Yoav Benjamini and Yosef Hochberg. Controlling the false discovery rate: a practical and powerful approach to multiple testing. *Journal of the Royal statistical society: series B (Methodological)*, 57(1):289–300, 1995.
